# Repurposing clofarabine against enteric pathogens via metabolic starvation

**DOI:** 10.64898/2026.10.09.757903

**Authors:** ND Deans, KL Naylor, S Srour, B Mallick, TK Pham, JW Sheffield, O Ansell, CD Edington-siddon, EA Burt, GP Stafford, MO Collins, D Humphreys

## Abstract

Enteropathogenic and enterohaemorrhagic *Escherichia coli* (EPEC/EHEC) cause severe global diarrhoeal disease, exacerbated by rising antimicrobial resistance. Pathogenesis relies on ‘attaching and effacing’ lesions, where bacteria generate actin pedestals to adhere, replicate, and destroy intestinal microvilli. To identify novel anti-virulence candidates targeting this process, we conducted a high-throughput phenotypic screen quantifying pedestal formation via high-content fluorescence microscopy. We identified the anticancer nucleotide analogue clofarabine as a potent inhibitor of EPEC growth and pathogenesis. Distinct from its oncology mechanism - inducing DNA chain termination and apoptosis - clofarabine competes with bacterial ATP-dependent processes to impede gene transcription, protein synthesis, and Type III Secretion System (T3SS) function. Remarkably, EPEC counteracts this inhibition by cleaving clofarabine, enabling restored protein synthesis and resumed growth. Furthermore, we report for the first time that clofarabine suppresses protein translation in human monocytes, suggesting an uncharacterised translational impact on mammalian cells. These findings reveal host and bacterial translation as interconnected vulnerabilities to purine antimetabolites, offering a novel strategy to disarm enteric pathogens while uncovering a bacterial enzymatic mechanism for neutralizing host-targeted therapeutics.

## Introduction

EPEC/EHEC remain major drivers of severe diarrheal disease and infant mortality [1, 2]. The global spread of antimicrobial-resistant (AMR) *E. coli* strains across more than 76 countries [3] led the World Health Organisation (WHO) to designate AMR *E. coli* a critical priority pathogen [4]. Despite this mounting threat, traditional antibiotic discovery offers low financial returns for pharmaceutical developers, creating a stagnant pipeline. Indeed, the number of newly approved antimicrobials has decreased significantly over recent decades and there is a major need for novel approaches to help discover potential antimicrobials [5]. Drug repurposing has thus emerged as an essential alternative strategy, leveraging existing clinical compounds to fast-track effective anti-virulence treatments [6]. This includes INP0341 and coumarin that have been successfully repurposed to inhibit Type III Secretion Systems (T3SS) and block effector delivery in Gram-negative pathogens [7, 8]. Accelerating these non-traditional approaches is critical to mitigate the escalating AMR crisis, which is projected to cause up to 10 million deaths annually by 2050 [9].

To cause disease, EPEC/EHEC infect intestinal epithelial cells by colonising the cell surface and inducing ‘attachment and effacement’ lesions characterised by the destruction of brush border microvilli [10]. A hallmark feature of the lesions is the formation of actin pedestals upon which EPEC/EHEC adhere, which establishes cell surface colonies of bacteria [11]. To generate actin pedestals, EPEC/EHEC assemble a T3SS, which is expressed via a virulence plasmid termed the locus of enterocyte effacement (LEE) that also encodes translocon components EspB/D [10, 12]. The T3SS injects LEE-encoded virulence effector proteins through the EspB/D translocon into the host cell that hijack the actin cytoskeleton. Underlying pedestal assembly is an effector known as the Translocated intimin receptor (Tir), which is delivered into the plasma membrane where it acts as a receptor for the bacterial outer membrane protein Intimin on the surface of EPEC [13]. Tir-Intimin interaction triggers host cell mediated tyrosine phosphorylation of Tir, which recruits the host adaptor protein Nck [14]. Once recruited, Nck activates Neural Wiskott-Aldrich syndrome protein (N-WASP) and the Actin-related Protein 2/3 (Arp2/3) complex, which drives actin polymerisation beneath adherent EPEC resulting in a signature actin pedestal [14]. EPEC/EHEC pedestal formation enables the pathogen to replicate upon its established niche and form micro-colonies of bacteria [15], which is driven by continued translocation of LEE- and non-LEE (Nle)-encoded effectors resulting in severe diarrheal disease [16, 17].

To identify novel anti-virulence candidates capable of disrupting host colonisation, we conducted a high-throughput phenotypic screen of a chemical library using high-content fluorescence microscopy to quantify actin pedestal formation. Strikingly, this screen revealed the anticancer nucleoside analogue clofarabine as the sole compound that effectively impeded EPEC infection. Clofarabine is traditionally used in oncology to induce DNA damage and apoptosis in host cells, therefore, its role in suppressing bacterial infection was unexpected. In this study, we set out to explore how this cancer therapeutic blocks EPEC pathogenesis and to investigate the molecular mechanisms governing its antibacterial activity.

## Results

### Establishing a High-throughput Imaging Approach to Assay EPEC Pedestal Formation

To identify novel inhibitors of EPEC pathogenesis, we created an automated, high content imaging pipeline to quantify actin-pedestal formation in host cells. Human Fibroblast HT1080 cells were infected with wild-type EPEC O127:H6 E2348/69 for 3 hours prior to fixation and imaging using the ImageXpress high-content fluorescence microscope (**Figure S1**). To monitor infection, we established a 384-well plate assay that combines immunofluorescent anti-Intimin detection of EPEC with phalloidin-FITC488 staining of actin pedestals and DAPI labelling of host cell DNA. This approach allowed unambiguous identification of adherent EPEC associated with actin pedestals (**Figure 1A, left: yellow arrow**).

**Figure 1:**
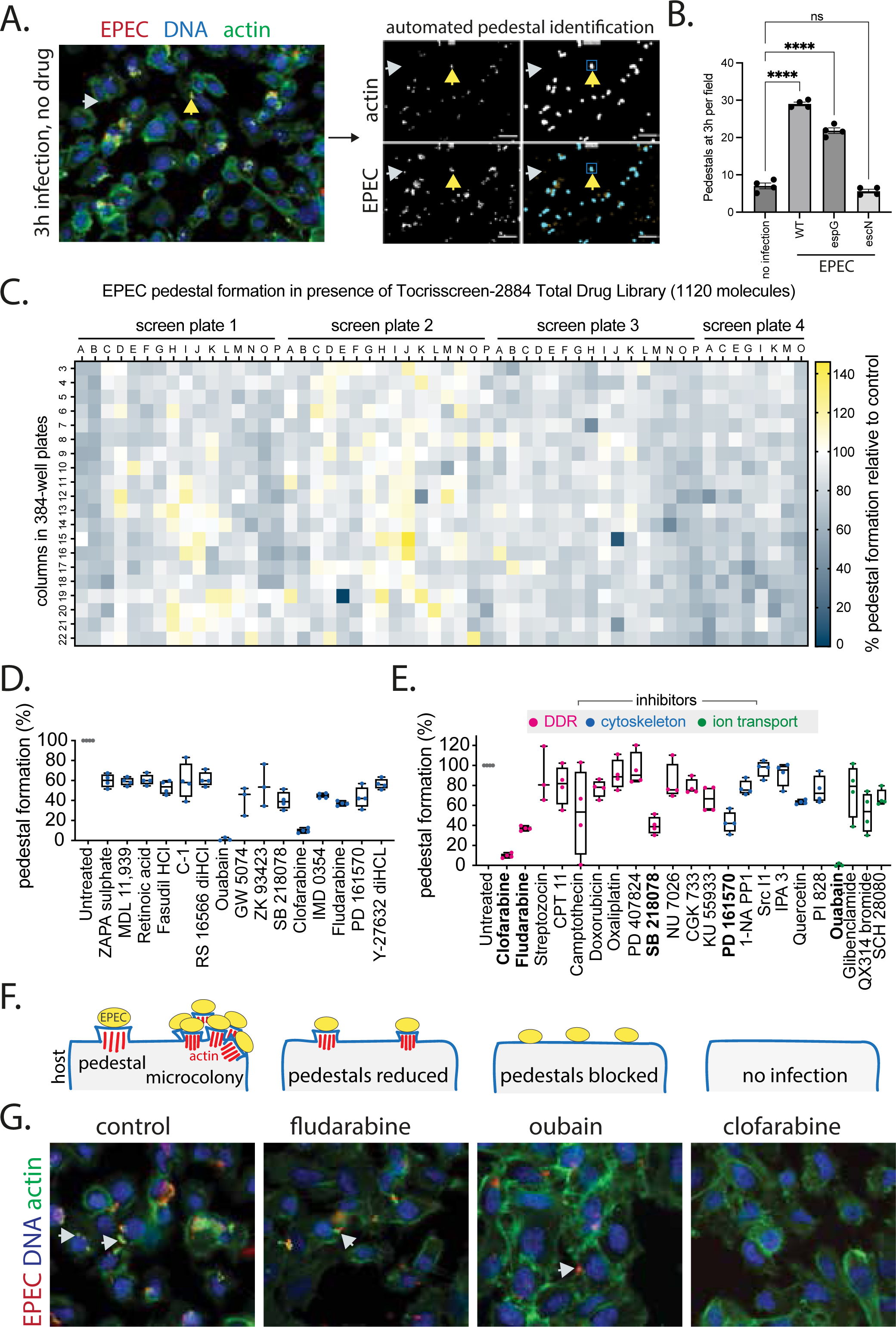
Clofarabine is a putative anti-infective agent. **(A)** Immunofluorescence imaging of HT1080 cells infected with (WT) EPEC for 3 hours prior to fixation, stained with phalloidin-488 for actin (green), anti-intimin for EPEC (red) and DAPI for nuclei (blue). Meta-Xpress CME software pipeline identifies actin foci and detects colocalised EPEC to define pedestals by colocalisation (yellow arrowhead); EPEC lacking actin colocalisation is highlighted by grey arrowhead. **(B)** Quantification of pedestals per field of view using Meta-Xpress data in HT1080 cells across untreated, WT EPEC, *ΔespG*, and *ΔescN* conditions. Statistical significance was determined by a one-way ANOVA with data presented as mean ±SEM ****p<0.0001. Individual circles represent fields of view from 3 independent experiments. **(C)** Heatmap showing the effect of Tociscreen total #2884 (Catalog no. 2884) drug library on EPEC actin pedestal formation in HT1080 cells relative to untreated control. Cells were seeded in 384-well plates treated with control DMSO (1%) or drugs solubilised in DMSO at 10μM prior to EPEC infection. Each compound was tested in a total of 4 technical repeats in 2 independent experiments. **(D)** Quantification of the 15 most effective inhibitors of EPEC pedestal formation with inhibition >40% identified in the drug screen in (C). **(E)** Quantification of drugs known to target DDR (magenta), cytoskeleton (blue) and ion transport (green) in EPEC, drugs highlighted in bold are reported in (D). Circles represent technical replicates from 2 biological replicates. **(F)** Schematic of infection phenotypes observed during screening of the Tocriscreen Total Library in control, Fludarabine, Oubain and Clofarabine (respectively, left-to-right). **(G)** Immunofluorescence imaging of HT1080 cells infected with WT EPEC post treatment with DMSO, Fludarabine, Oubain and Clofarabine, respectively, stained with phalloidin-488 for actin (green), anti-intimin for EPEC (red) and DAPI for nuclei (blue); white arrowheads highlight pedestals.

To automate pedestal quantification and distinguish pathogen-induced structures from host cytoskeletal remodelling, we configured MetaXpress Custom Module Editor (CME) software to detect actin foci that co-localise with EPEC (**Figure 1A**, **right**: **yellow arrowhead**). The pipeline successfully discriminated between non-pedestal-associated, adherent EPEC (**grey arrowhead**) and pedestal-inducing EPEC (**yellow arrowhead**). Because pedestal orientation varies relative to the bacterial body, the software expanded the region of interest surrounding each actin focus to capture associated bacteria, quantifying EPEC microcolonies across nine fields of view per well (**Figure 1A**, **right**: **blue square**).

To evaluate assay stringency, we compared pedestal induction between wild-type (WT) EPEC and the T3SS-deficient mutant *ΔescN* (Gauthier et al 2003) (**Figure 1B**). The automated pipeline detected ∼30 pedestals per field of view for WT EPEC, whereas *ΔescN*-infected cells yielded ∼5 counts per field. Because *ΔescN* cannot induce pedestals, these low-level signals represent baseline assay noise (pseudo-pedestals), consistent with uninfected controls (∼6 background counts). Additionally, infecting cells with a mutant lacking EspG (*ΔespG*) - a non-essential effector known to promote efficient pedestal assembly [18] - resulted in an intermediate, moderate reduction in pedestal counts (∼20 per field) (**Figure 1B**). Together, these data validate a sensitive, automated platform for quantifying EPEC actin pedestal formation.

### Identification of Clofarabine as an Inhibitor of EPEC Pedestal Formation

To identify potential anti-infectives targeting EPEC pedestal formation, we screened the Tocriscreen-2884 Total Compound Library (1,120 compounds) across four 384-well plates **(Table S1)** during a 3-hour infection (**Figure 1C, S1**). Screening heatmaps revealed that compounds either enhanced pedestal formation (110%–140%; yellow) or modestly suppressed it (40%–100%; grey) (**Figure 1C**). Relative to infection of DMSO-treated control cells, 15 compounds reduced pedestal formation by >40%, with the most potent inhibition observed for ouabain and clofarabine (<10% relative to control) (**Figure 1D, S2–S5**). Quantification of host cell nuclei confirmed that none of these 15 compounds reduced host cell numbers, indicating that the observed anti-virulence phenotypes were due to the specific inhibition of EPEC infection rather than host cytotoxicity **(Figure S6A).**

Clofarabine (Evoltra) is a second-generation purine nucleoside analogue that activates host DNA damage responses (DDR) and apoptosis in cancer cells [19], whereas ouabain is a Na+/K+-ATPase inhibitor with broad therapeutic applications [20, 21]. Although we initially hypothesised that direct cytoskeletal inhibitors would disrupt pedestal formation, none of the molecules tested in the category attenuated pedestal formation (**Figure 1E, blue**). Similarly, among the ion transport inhibitors in the library, only ouabain attenuated pedestal assembly (**Figure 1E, green**). Finally, while diverse DDR activators and regulators were present in the library, only clofarabine, fludarabine, SB218078 (a Chk1 inhibitor) and PD161570 (inhibitor of fibroblast growth factor receptor 1 (FGFR1)) suppressed pedestal formation by >40% (**Figure 1E, magenta**). Fludarabine is a structurally related purine nucleoside analogue that served as a key predecessor to clofarabine, which was subsequently developed to improve metabolic stability and therapeutic efficacy [19].

Analysis of the fluorescence images from the screen revealed distinct infection phenotypes, as schematised in **Figure 1F** and shown in **Figure 1G**. Among the 15 hits, 13 compounds permitted bacterial adhesion while reducing pedestal formation, exemplified by fludarabine (**Figure 1G, fludarabine**). A second phenotype was observed with ouabain, which completely abrogated pedestal formation while still allowing bacterial colonisation and therefore infection (**Figure 1G, ouabain**). Finally, treatment with clofarabine resulted in a complete loss of both bacterial adhesion and pedestal formation (**Figure 1G, clofarabine**). Together, these findings identify clofarabine as the sole compound in the library capable of completely blocking EPEC infection, highlighting its potential as a candidate for drug repurposing.

### Clofarabine Exhibits Selective Antimicrobial Activity Against Bacteria

Having identified clofarabine as a hit in pedestal inhibition assays, we evaluated whether its phenotypic effect is due to broad antibacterial growth suppression or strain-specific activity (**Figure 2**). In short-term growth assays (4.5 h), clofarabine strongly suppressed EPEC growth (*p* < 0.0058 relative to untreated controls), maintaining OD600 near baseline levels comparable to the bactericidal control, kanamycin (**Figure 2A**). However, extended 24-hour monitoring revealed that clofarabine delays EPEC exponential entry by approximately 6–7 hours compared to untreated controls before cells eventually reach stationary phase at 11–12 hours (**Figure 2B**). This demonstrates a transient bacteriostatic effect rather than the sustained bacterial clearance seen with kanamycin. (**Figure 2B**).

**Figure 2:**
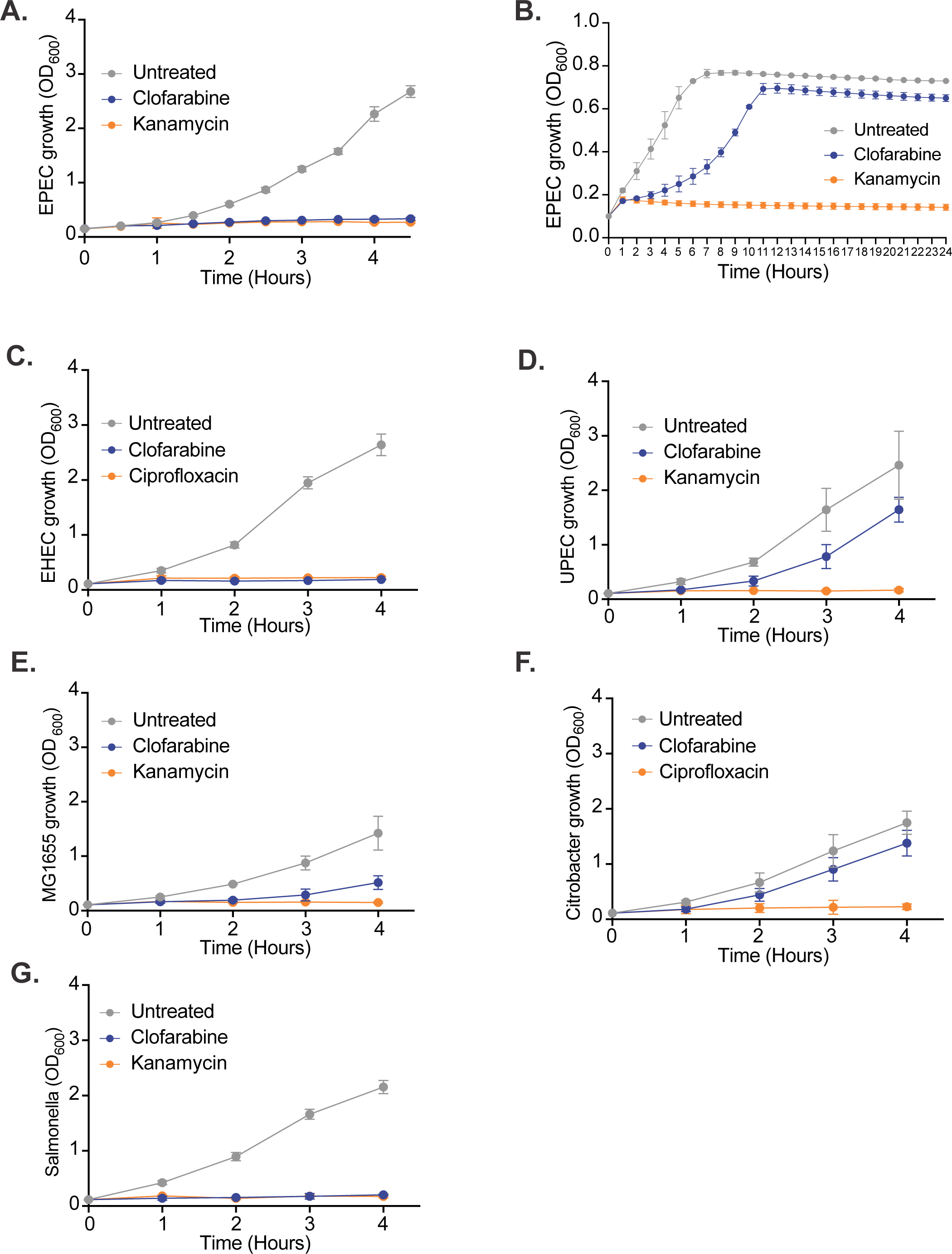
Clofarabine inhibits EPEC and EHEC growth. Growth curves of various bacterial stains starting at an initial OD600 of 0.1. Bacteria were cultured in DMEM and treated with 10μM Clofarabine (blue), a positive control antibiotic (kanamycin or ciprofloxacin; orange), or left untreated (grey). **(A)** EPEC growth over 4.5 hours, **(B**) EPEC growth over 24 hours, **(C)** EHEC, (**D)** UPEC, **(E)** MG1655, **(F)** Citrobacter, and **(G)** Salmonella Typhimurium.

To determine whether clofarabine sensitivity correlates with T3SS-dependent pathogenesis, specific metabolic features, or evolutionary lineage, we tested a panel of enteric bacteria spanning LEE-positive pathogens, non-LEE T3SS pathogens, and non-T3SS controls (**Figure 2C-2G**). Enterohemorrhagic *E. coli* (EHEC), an A/E pathogen sharing the LEE-encoded T3SS machinery with EPEC, was completely growth-blocked over 4 hours (*p* < 0.045), mirroring the bactericidal ciprofloxacin control (*p* = 0.0451; **Figure 2C**). In contrast, non-LEE extraintestinal strains - uropathogenic *E. coli* (UPEC) and non-pathogenic *E. coli* K-12 (MG1655) - exhibited only non-significant partial growth attenuation characterised by a reduced exponential slope rather than complete baseline suppression (*p* = 0.5139 and *p=* 0.1534, respectively; **Figure 2D, 2E**).

To test whether LEE expression drives sensitivity, we assayed *Citrobacter rodentium*, a murine A/E pathogen encoding a LEE island homologous to EPEC and EHEC. Surprisingly, *C. rodentium* was resistant to clofarabine, displaying growth kinetics similar to untreated controls (*p*=0.0843; **Figure 2F**). Finally, we evaluated *Salmonella* Typhimurium, an enteropathogen that lacks LEE but expresses T3SS-1 from *Salmonella* pathogenicity island 1 (SPI-1) *in vitro*. Clofarabine suppressed *S.* Typhimurium growth (*p* < 0.025), matching the acute sensitivity observed in EPEC and EHEC (*p=*0.0256; **Figure 2G**). Since growth inhibition occurs independently of LEE carriage and sensitivity varies among T3SS-expressing species, clofarabine uptake and activity do not depend on the LEE needle complex, highlighting selective bacteriostatic susceptibility among specific enteric pathogens.

### Clofarabine Induces Transcriptomic Changes Suppressing Virulence and Alters Purine Metabolism

The growth suppression observed in enterobacteria, EPEC, raises the question of how clofarabine affects the bacterial transcriptome. To understand this, we performed RNA-seq transcriptomic profiling on EPEC under treated (10 µM clofarabine; CIF) and untreated conditions across two growth phases: at OD600 =0.2, when growth is arrested, and at OD600 =1.0, when EPEC have initiated recovery (**Figure 2B**, **3**). Principle component analysis (PCA) showed clear separation of the expression profiles by treatment PC1 (43.08% of variance), with clofarabine-treated samples at OD600 = 0.2 clustering distinctly from untreated controls (**Figure 3A**). Interestingly, while clofarabine-treated samples at OD600 =1.0 also separate from their corresponding untreated controls along PC2 (15.27% of variance), they cluster much closer to the control group along PC1 than the OD600 =0.2 treated samples. A hierarchical clustering heatmap of overall gene expression confirms the broad transcriptional shift induced by clofarabine at OD600 =0.2, whereas the expression profile at OD600 =1.0 shows more modest differences between treated and untreated conditions (**Figure 3B**). These data indicate that clofarabine’s primary transcriptomic impact occurs during early growth arrest, with gene expression relaxing back toward baseline as cells overcome bacteriostasis.

**Figure 3:**
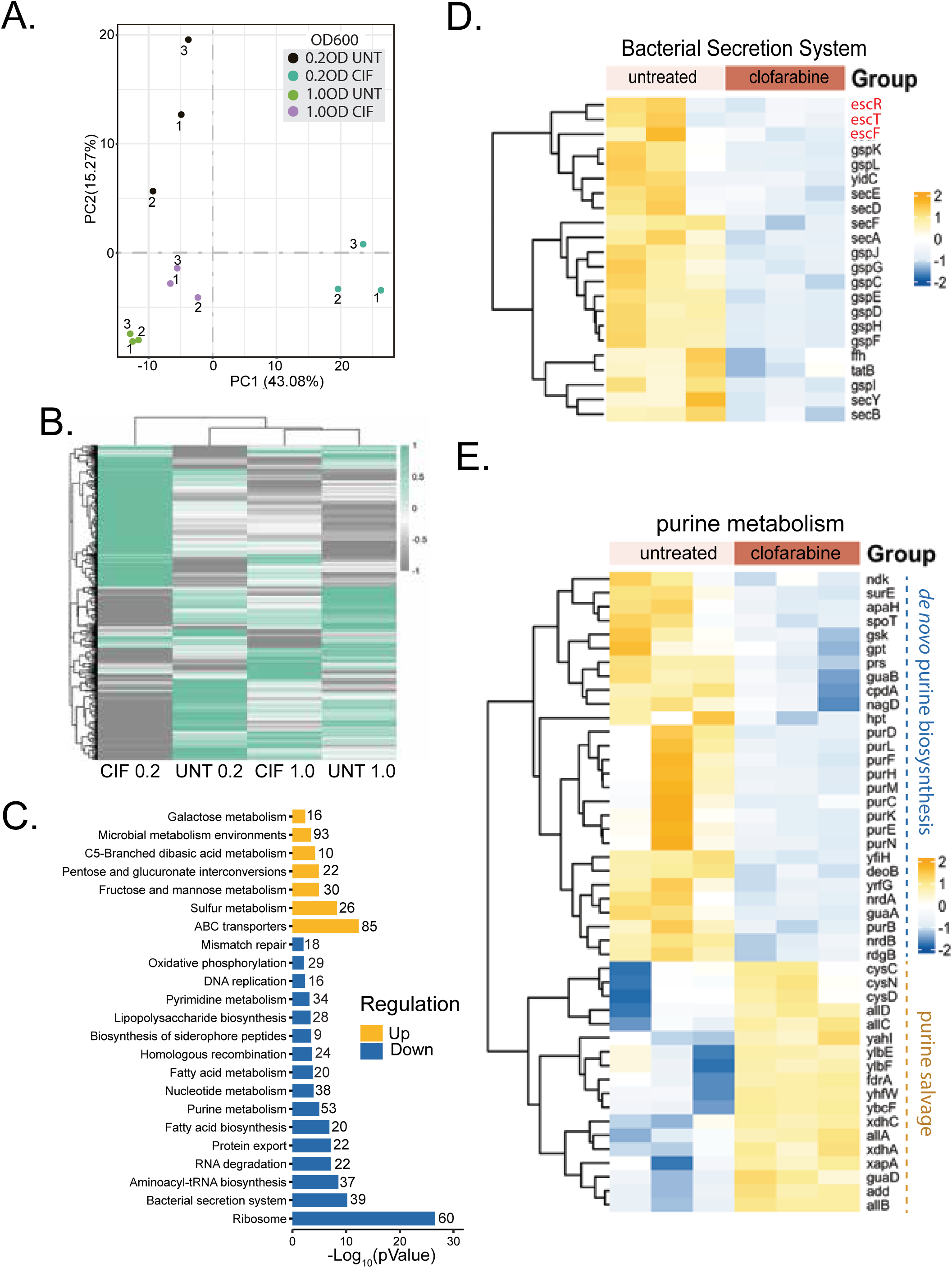
Clofarabine inhibits EPEC expression of ribosomal and virulence genes. **(A)** Principal component analysis (PCA) of transcriptomes from untreated (UNT) or CIF-treated (CIF) EPEC at OD600 0.2 (early log phase; UNT 0.2 and CIF 0.2) or EPEC at OD600 1.0 (late log phase; UNT 1.0 and CIF 1.0). **(B)** Hierarchical clustering heatmap of the 3,838 differentially expressed genes (DESeq2 Padj ≤0.05 in at least one pairwise comparison) across untreated and clofarabine-treated EPEC cultures at OD_600_ 0.2 and 1.0. Data are represented as row-scaled log_2_(FPKM+1) values using the mean of three biological replicates. Genes and conditions were clustered using Euclidean distance and complete linkage. **(C)** KEGG Pathway analysis comparing CIF vs UNT of OD600 0.2. Values indicate total genes enriched in the pathway. **(D)** Heatmaps showing the relative expression of significant genes bacterial secretion systems with T3SS/LEE genes in red, and **(E)** significant genes involved in purine metabolism with relative changes in *de novo* purine biosynthesis and purine salvage indicated, right. Values in (D), (E) are mapped with yellow showing up-regulated genes and blue showing down-regulated genes.

KEGG pathway enrichment analysis of differentially expressed genes at OD600 =0.2 revealed widespread transcriptional reprogramming upon clofarabine treatment (**Figure 3C**). Notably, clofarabine led to a strong downregulation of 60 genes encoding the ribosome - comprising 54 ribosomal proteins across the 30S (21 proteins) and 50S (33 proteins) subunits [22, 23]- as well as 37 genes governing aminoacyl-tRNA biosynthesis (**Figure 3C, S3B**). We also observed suppression of bacterial secretion system genes (39 downregulated genes), including structural needle apparatus components (*escR*, *escT*, *escF*; highlighted in red) and general secretion pathway factors (*secA*, *secD*, *secE*, *secF*) (**Figure 3C**, **3D**). This dual repression provides a clear rationale for both the bacteriostatic growth arrest (**Figure 2**) and the loss of actin pedestal formation (**Figure 1**).

Furthermore, clofarabine treatment triggered a striking metabolic shift in purine pathways, characterised by the upregulation of purine salvage genes alongside a significant downregulation of genes governing *de novo* purine biosynthesis (**Figure 3C**, **3D**). Specifically, key enzymes in the *de novo* purine biosynthetic pathway - including the *pur* operon cluster (*purD*, *purL*, *purF*, *purH*, *purM*, *purC*, *purK*, *purE*, and *purN*) - were repressed (**Figure 3D**), consistent with severe purine nucleotide disruption induced by this adenine nucleoside analogue. Interestingly, expression of SOS response genes showed no significant change (**Figure 3D**). To confirm this at the protein level, immunoblotting demonstrated that clofarabine treatment failed to induce RecA expression, which supports that there is a global translational arrest rather than an active SOS response (**Figure S3C**). Together, these results demonstrate that clofarabine acts via a dual mechanism: disrupting purine metabolism while simultaneously suppressing ribosomal translation and T3SS virulence machinery.

### Clofarabine Inhibits the Function of the EPEC Type 3 Secretion System

Given the observed reduction in expression of bacterial secretion system genes (**Figure 3D**), we next evaluated whether clofarabine inhibits T3SS directly during infection (**Figure 4**). To establish dose-dependency, HT1080 cells were infected with EPEC (MOI = 50) across a clofarabine titration (0.1 µM to 10 µM), alongside uninfected, vehicle (DMSO), and antibiotic (kanamycin) controls (**Figure 4A**). Confocal microscopy demonstrated robust actin pedestals in DMSO-treated controls, characterised by localised recruitment of host F-actin (red) directly beneath attaching bacteria (intimin, green; host/bacterial nuclei, DAPI, blue). At 10 µM and 5 µM, clofarabine completely abolished pedestals, phenocopying the uninfected and kanamycin-treated controls (**Figure 4A**). Stepwise reduction of clofarabine (1 µM to 0.1 µM) resulted in a clear, dose-dependent restoration of bacterial attachment and host F-actin recruitment into pedestals (**Figure 4A**). Quantification confirmed a significant reduction in pedestal-positive cells across 0.2 µM, 0.5 µM, 1 µM, 5 µM, and 10 µM clofarabine compared to DMSO controls (**Figure 4B**).

**Figure 4:**
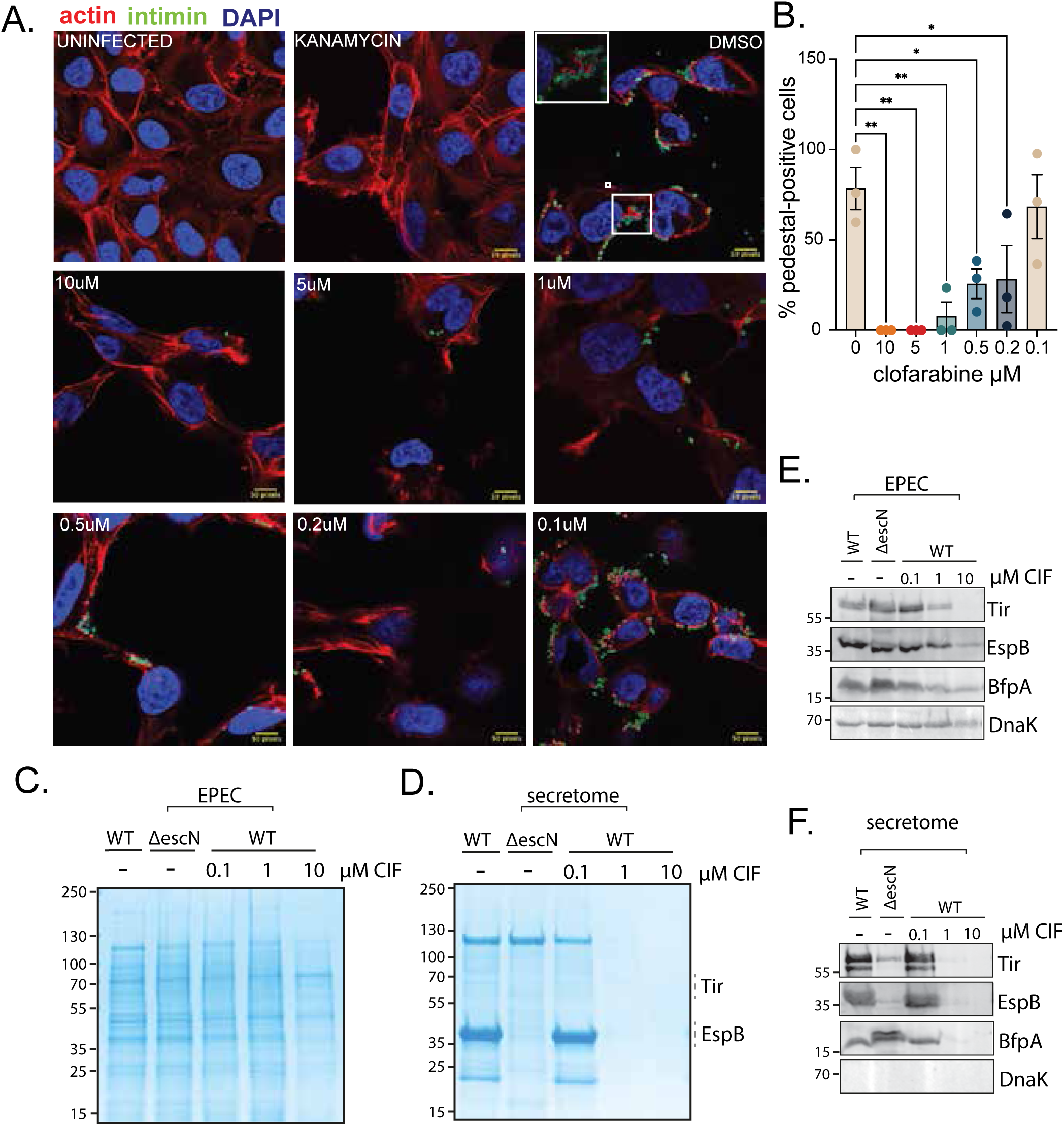
Clofarabine inhibits the function of the EPEC T3SS. **(A)** Immunofluorescence imaging of HT1080 cells infected with WT EPEC incubated with varying concentrations of Clofarabine (10μM, 5μM, 1μM, 0.5μM, 0.2μM and 0.1μM), kanamycin or DMSO compared to the uninfected control. Cells are stained with phalloidin-568 (red) anti-intimin for EPEC (green) and DAPI for nuclei (blue). **(B)** Quantification of pedestals per field of view. **(C)** Coomassie blue-stained SDS-PAGE gel showing total EPEC whole-cell lysate from WT EPEC, ΔescN (T3SS-deficient mutant) and WT EPEC treated with increasing concentrations of Clofarabine (0.1,1.10μM CIF). **(D)** Coomassie blue-stained SDS-PAGE gel of secreted proteins (secretome) collected from the supernatant of the indicated EPEC strains and treatment conditions. Hashed lines indicated where gel slices (lane 1: WT, -) were excised for LC-MS/MS analysis confirming presence of Tir and EspB as labelled, right. **(E)** Immunoblot analysis of samples from (C) using indicated antibodies against effector (Tir, EspB), bundle forming pili protein BfpA and DnaK (loading control). **(F)** Immunoblot analysis of samples from (D) using indicated antibodies. Molecular weight markers in kDa indicated left in (C), (D), (E), and (F).

To determine whether the loss of pedestal assembly stems from suppressed effector expression or defective protein secretion, we analysed whole-cell lysates (**Figure 4C**, **4E**) and filtered culture supernatants (secretome) (**Figure 4D**, **4F**) from WT EPEC and a T3SS-null mutant (*ΔescN*). SDS-PAGE with Coomassie blue staining of whole-cell lysates demonstrated that total intracellular protein profiles remained broadly comparable across treated and untreated conditions (**Figure 4C**). To identify and quantify specific proteins in the culture supernatant, bands from the secretome samples separated by SDS-PAGE were excised and analysed by mass spectrometry **(Table S2)** confirming the presence of key T3SS effectors, EspB and Tir. However, targeted immunoblotting of whole-cell lysates revealed selective alterations in intracellular protein pools (**Figure 4E**). While the cytosolic control DnaK and the T3SS translocator EspB remained stable across conditions, intracellular Tir expression was selectively depleted at 10 µM clofarabine (**Figure 4E**). In the secretome, 1 µM and 10 µM clofarabine completely eliminated secretion of the essential pore-forming translocator EspB and the actin-pedestal receptor Tir, while DnaK (a cytoplasmic protein), was undetected. (**Figure 4F**).

Finally, we evaluated the bundle-forming pilus subunit BfpA—a primary EPEC adhesin encoded on the pEAF plasmid that relies on the general secretion (Sec) pathway (*secY*, *secB*) for inner membrane translocation [24]. While intracellular BfpA synthesis was moderately attenuated at 10 µM clofarabine in whole-cell lysates (**Figure 4E**), its extracellular release into the secretome was completely abolished at 1 µM and 10 µM (**Figure 4F**). This selective loss of extracellular BfpA indicates that while precursor synthesis persists, sec-dependent inner membrane translocation and subsequent pilus assembly are blocked, aligning with the transcriptomic downregulation of sec machinery observed in Figure 3D.

Together, these results demonstrate that clofarabine disarms EPEC pathogenicity through a dual mechanism: blocking functional secretion of T3SS translocators (EspB) and effectors (Tir) to prevent host pedestal formation, while simultaneously impairing Sec-dependent export of major pilus adhesins (BfpA).

### Clofarabine Suppresses Ribosomal Expression and Protein Synthesis

Our functional assays established that clofarabine suppresses T3SS effector expression and secretion thereby disarming EPEC virulence (**Figure 4**), but together with the simultaneous growth delay (**Figure 2**), the data pointed toward a shutdown of protein synthesis. Since transcriptomic profiling highlighted downregulation of ribosomal gene clusters (**Figure 3C**), we investigated whether clofarabine directly impairs ribosomal gene expression and translation efficiency (**Figure 5**). Transcriptomic analysis of ribosomal operons showed global suppression of ribosomal gene expression in clofarabine-treated samples, compared to untreated controls (**Figure 5A**). Hierarchical clustering confirmed that all 30S and 50S ribosomal structural genes were down-regulated, suggesting a transcript-level shutdown of bacterial ribosome biogenesis machinery.

**Figure 5:**
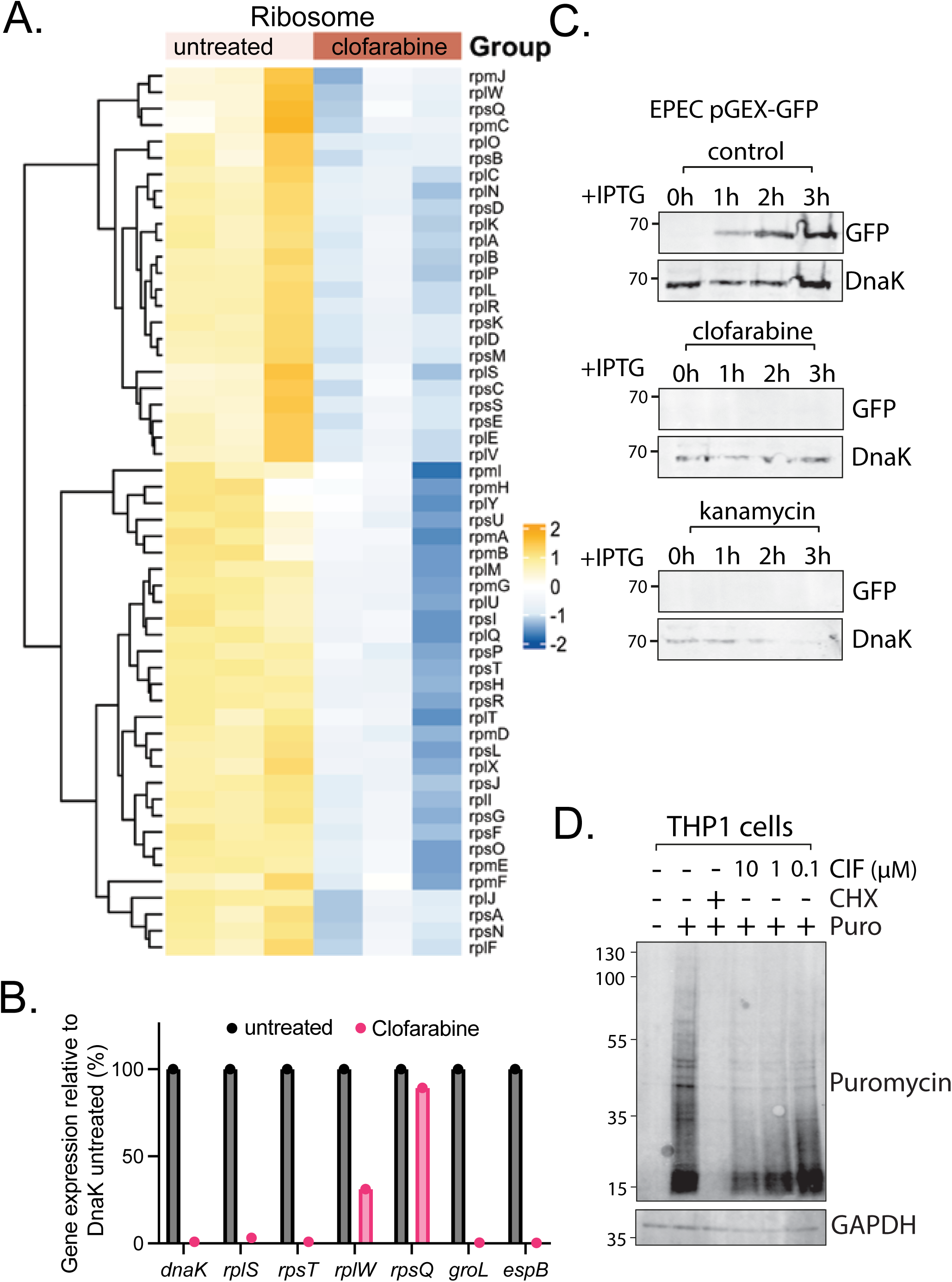
Clofarabine suppresses bacterial ribosomal machinery and shuts down protein synthesis. (**A**) Hierarchical clustering heatmap showing relative transcript expression levels of ribosomal genes in untreated versus clofarabine-treated EPEC at OD600 =0.2. Upregulated genes highlighted in yellow and downregulated in blue. (**B**) qRT-PCR analysis of relative gene expression levels (%) in untreated (black bars) and clofarabine-treated (magenta bars) EPEC. Expression target ribosomal genes (*rplS, rest, rplW, rpsQ*), molecular chaperones (*dnaK, groL*), and virulence translocators (*espB*) were normalised against the untreated control relative to the *dnaK* internal reference. Data are presented as mean+/-SEM, n=1. (**C**) Immunoblot analysis of *de novo* protein synthesis using an IPTG-inducible pGEX-GFP EPEC reporter strain. GFP was induced with IPTG (0.5mM) and assayed at 1hr intervals over 3 hrs under control, clofarabine (10μM) or kanamycin (50μg/ml). GFP reporter levels and DnaK loading controls were assayed to show translational arrest. (D) Representative western blot of a SUnSET assay measuring active protein sysnthesis in THP1 monocytes. Cells were treated with decreasing concentrations of clofarabine (left-to-right; 10, 1 and 0.1μM), cycloheximide (CHX; positive control, 100μg/ml), or vehicle control before puromycin pulse-labeling. Immunoblotting was performed with anti-puromycin antibodies; GAPDH serves as a loading control.

To further validate the transcriptomic findings, we performed targeted quantitative real-time PCR (qRT-PCR) on key components normalised against the *dnaK* internal control (**Figure 5B**). Standard qRT-PCR normalisation relies on house-keeping control genes such as *dnaK*. However, because clofarabine induces widespread transcriptional shutdown that suppresses endogenous *dnaK* expression, normalising treated samples directly against their internal *dnaK* levels generates an artifactual apparent upregulation of target transcripts (**Figure S7C**). To accurately quantify relative transcript loss without this confounding artifact, gene expression in both untreated and clofarabine-treated samples was normalised against the baseline *dnaK* expression of the untreated control (**Figure 5B**). Clofarabine treatment (magenta bars) led to a near-complete loss of transcripts encoding core ribosomal proteins (*rplS*, *rpsT*, *rplW*), the chaperones *groEL, dnaK*, and the translocon *espB* relative to untreated controls (black bars). Interestingly, *rpsQ* (encoding 30S ribosomal protein S17) was spared from this shutdown, retaining ∼90% of baseline transcript levels and highlighting specific transcript sensitivity within ribosomal genes.

To test whether this transcriptomic loss leads to translational arrest, we monitored *de novo* protein synthesis using an IPTG-inducible reporter system (EPEC pGEX-GFP) (**Figure 5C**). In this system, GFP expression is dependent on the activation of an IPTG-inducible promoter. Upon IPTG induction, untreated control cells showed accumulation of GFP in a time dependent manner over a 3hr period (**Figure 5C**). However, with clofarabine-treatment GFP expression was abolished, despite the presence of IPTG. This mimics the total blockade of GFP expression seen with the translational inhibiting antibiotic kanamycin (**Figure 5C**).

Finally, to assess whether this protein synthesis defects extends to eukaryotic host cells, we measured protein synthesis in human macrophage-like THP1 cells using SUnSET puromycin incorporation assay **(Figure 5D)**. While the positive control cycloheximide (CHX) completely blocked translation, highlighted by the loss of puromycin, clofarabine (ClF) treatment caused a dose dependent reduction in puromycin incorporation (0.1 – 1μM) relative to untreated control. Together, these data suggest that ClF inhibits protein synthesis at the transcriptional level in the host cell, meanwhile repressing ribosomal gene expression in bacteria driving a global arrest in active bacterial translation and attenuating translation in mammalian cells.

### Adenosine Rescue and Bacterial Degradation of Clofarabine Reverse Toxicity

Clofarabine is a structural purine nucleoside analogue. This led us to hypothesise that the translational arrest observed and T3SS blockade stems from competitive incorporation of clofarabine into nucleotide metabolic pathways, effectively inducing a state of purine starvation. To test whether exogenous purines could outcompete clofarabine and rescue bacterial physiology, we supplemented EPEC with increasing concentrations of adenosine (**Figure 6A-D**). While clofarabine alone (10μM) maintained a robust suppression of growth over 4 hours (**Figure 6A**: *p* < 0.02 relative to untreated control), co-treatment with adenosine restored EPEC growth kinetics in a dose-dependent manner (**Figure 6B-D**). Relative to controls, 0.1mM (**Figure 6B**: *p* < 0.0001) and 1mM adenosine (**Figure 6C**: *p* < 0.014), clofarabine still significantly suppressed growth but at 10mM adenosine, the growth inhibitory effect of CIF was fully restored to untreated conditions (**Figure 6D**: *p* < ns).

**Figure 6:**
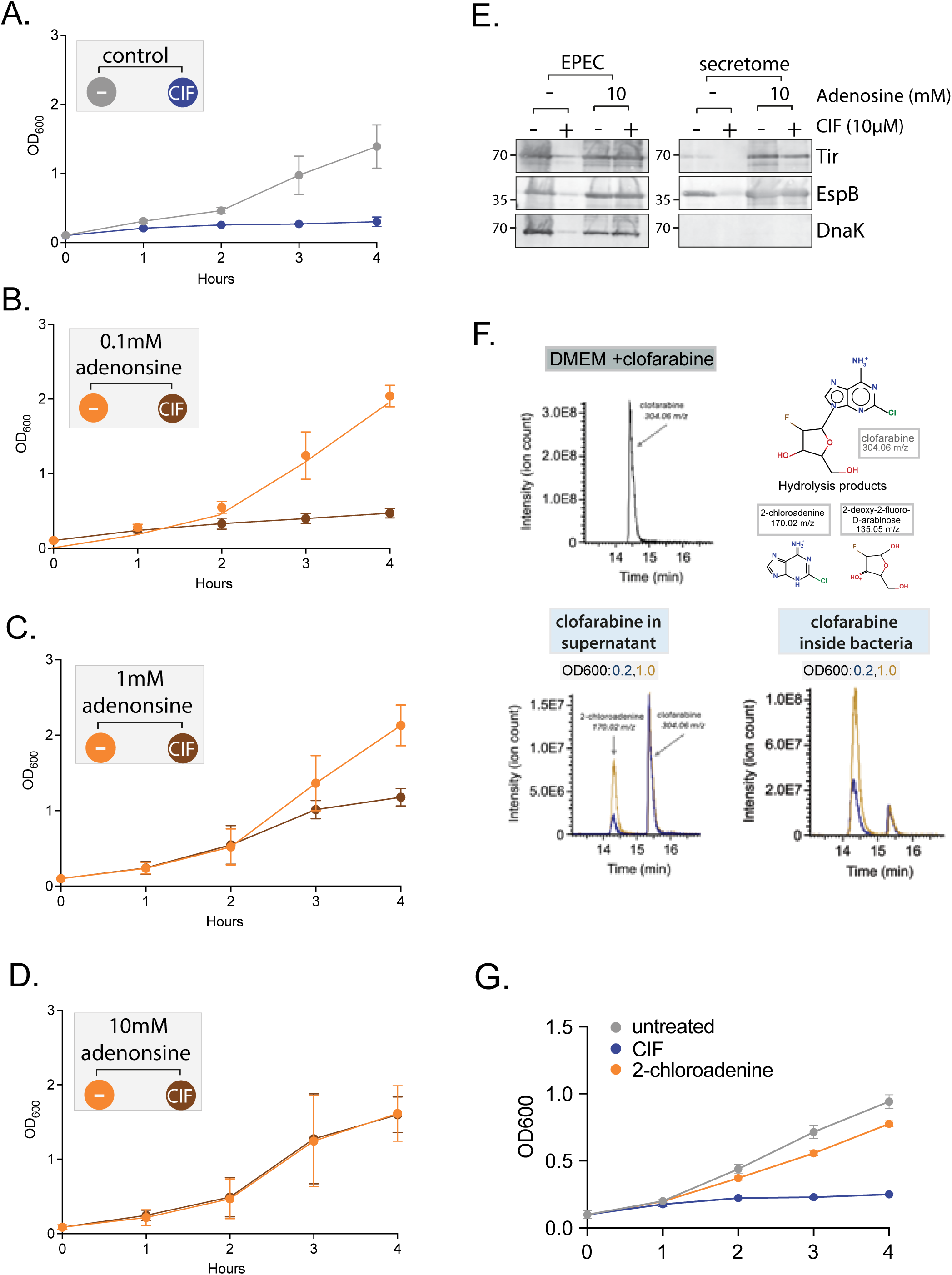
Competitive purine starvation drives clofarabine toxicity and is reversed by bacterial degradation of clofarabine. (**A-D**) Growth of EPEC (OD600) over 4 hours under specified treatment conditions demonstrating dose-dependent adenosine rescue. (**A**) Growth curve of untreated control versus clofarabine-treated EPEC (CIF; 10μM). (**B-D**) Co-treatment of CIF (10μM) with increasing concentrations of exogenous adenosine (**B**) 0.1mM (**C**) 1mM and (**D**) 10mM adenosine compared to adenosine-only controls. Data represent mean +/− SEM (n=3 independent experiments). (**E**) Immunoblot analysis of whole cell lysates (EPEC) and clarified secretome (secretome) probing Tir, EspB and DnaK loading control. Cultures were treated with (+) and without (−) clofarabine (CIF; 10μM) in the presence of exogenous adenosine (10mM) to assay T3SS and rescue. (**F**) Liquid chromatography-mass spectrometry (LC-MS) characterisation of clofarabine degradation and metabolic cleavage products. Top: LC-MS extracted ion chromatogram of parent clofarabine (304.06 m/z) in cell-free DMEM media (top left) alongside chemical structures and m/z values for intact clofarabine and its enzymatic hydrolysis products (top): 2-deoxy-2-fluoro-D-arabinose (right; 135.05 m/z) and 2-chloroadenine (left; 170.02 m/z). Bottom: Extracted ion chromatograms tracking clofarabine degradation at indicated OD600 readings with EPEC culture supernatants (left) and intracellular bacterial fraction (right), showing active drug internalisation and hydrolysis product accumulation (170.02 m/z). (n = 3) (**G**) Growth of EPEC (OD600) treated with vehicle control, clofarabine (CIF), or the cleaved base fragment 2-chloroadenine over 4 hours (n=3).

To determine if purine supplementation restores virulence expression, we blotted for virulence effectors, EspB and Tir in whole cell lysates (EPEC) and secretome fractions (**Figure 6E**). Adenosine at 10mM restored both intracellular accumulation and extracellular secretion of both T3SS effectors in the presence of clofarabine (**Figure 6E**). This competitive rescue demonstrates that clofarabine does not cause permanent structural damage to the T3SS needle, but rather induces a reversible metabolic block driven by purine depletion through competition. We reasoned that clofarabine-mediated suppression of EPEC may be influenced by the formulation of growth media. Consistent with this hypothesis, clofarabine inhibited expression of Tir and EspB in defined media lacking an exogenous source of purines such as DMEM or minimal media, but this was not the case in rich media such as LB where Tir and EspB were fully expressed (**Figure S7D**).

**Figure 2B** shows restoration of growth kinetics over a 24-hour period, suggesting the clofarabine effect to be transient in 6-8h of treatment. To uncover the mechanism behind this transient effect, we used liquid chromatography-mass spectrometry (LC-MS) to track drug stability in culture fractions (**Figure 6F**). Clofarabine remained intact (304.06 m/z) and stable in cell-free culture media (DMEM). However, when EPEC was cultured with clofarabine, extracellular and intracellular fractions showed that the active drug was cleaved into its enzymatic hydrolysis products, 2-cloroadenine (170.02 m/z) and 2-deoxy-2-fluoro-D-arabinose (135.05 m/z) (**Figure 6F**). Profiles of intracellular clofarabine show that EPEC actively internalises and degrades clofarabine at the 0.1 starting OD600 (**Figure 6F**).

Having established that EPEC cleaves clofarabine, we next tested whether this degradation product retained bacterial toxicity. To test this, we evaluated the effect of the primary cleaved base fragment 2-cloroadenine on bacterial growth (**Figure 6G**). 2-chloroadenine failed to suppress EPEC growth (*p* = 0.125 relative to untreated control), unlike the parent clofarabine (*p* = 0.015). Together these findings establish that clofarabine acts as a competitive purine antimetabolite that shuts down translation and T3SS function. This effect is not permanent and can be reversed. EPEC has a survival mechanism that internalises and metabolises clofarabine, clearing the active component into non-toxic products, which allows for restoration of bacterial translation and T3SS function.

## Discussion

Rapidly rising antimicrobial resistance pushes the need for non-traditional therapeutics that disarm virulence mechanisms without exerting selective pressure for resistance. In this study, we established an automated high-content imaging screen to identify small-molecule inhibitors of EPEC actin pedestal formation (**Figure 1**). We identified clofarabine, an FDA-approved purine nucleoside analogue, as a potent dual-action inhibitor that blocks T3SS machinery and halts acute bacterial growth (**Figures 1**, **2**). Through our transcriptomic profiling, quantitative protein assays, and metabolic rescue experiments, we demonstrate that clofarabine acts as a competitive purine antimetabolite. Clofarabine works by inducing nucleotide starvation, triggering global repression of ribosomal biogenesis and secretion machinery, culminating in translational arrest and T3SS inactivation (**Figures 3–5**). Meanwhile, EPEC counteracts this metabolic block through internalising and cleaving clofarabine into non-toxic sub-components to eventually restore growth and virulence (**Figure 6**).

A pivotal finding of our work is that clofarabine’s anti-virulence strategy is tightly coupled to a fundamental disruption of bacterial nucleotide metabolism. The EPEC T3SS is a high-demand biological machine requiring ATP consumption for protein translocation export, as well as protein translation to maintain the needle’s structure and effector assembly [25, 26]. Our transcriptomic and functional profiling revealed that clofarabine downregulates *de novo* purine synthesis operons, while upregulating purine salvage and catabolic pathways (**Figure 3D**). This striking metabolic shift is characteristic of purine starvation. This coordinated response suggests that under clofarabine-induced purine starvation, EPEC initiates a metabolic emergency response, shifting into an “energy-conserving” state. This is done by downregulating non-essential, energy-intensive processes, allowing the bacterium to prioritise metabolic survival over host colonisation. Interestingly, exogenous adenosine supplementation fully restored both EPEC growth and secretion of essential extracellular virulence effectors (Tir, EspB) in a dose-dependent manner (**Figure 6**). This competitive rescue further proves that clofarabine does not cause permanent damage to the T3SS, but rather enforces a reversible metabolic blockade, further supporting the transient “energy-conserving” state.

Even though clofarabine induces prominent translational arrest, interestingly, it does so through a mechanism different from classical DNA-damaging antibiotics. Fluoroquinolones, like ciprofloxacin, that cause DNA damage and translational arrest, trigger the bacterial SOS response, inducing RecA expression [27]. Although RecA helps the bacterium repair its genome, it comes at a cost, where it allows for horizontal gene transfer and resistance to certain antibiotics [28]. Clofarabine fails to induce RecA (**Figure S6C**), and there are no significant changes in the expression of SOS genes. This the conclusion that, in EPEC, clofarabine does not act as a genomic DNA-damaging agent that triggers a hazardous bacterial SOS emergency response. Instead, clofarabine drove a global downregulation of 30S and 50S ribosomal subunits and molecular chaperones, resulting in a translational blockade (**Figure 5**). This disruption of nucleotide metabolism without SOS induction is a major therapeutic advantage. It suggests that purine antimetabolites could be further explored as they can target A/E pathogens like EPEC/EHEC without risks of antibiotic-induced resistance.

Through our strain specificity panel, we provided key insights into the requirements of clofarabine susceptibility, uptake, and metabolic activation (**Figure 2**). Clofarabine suppressed LEE-positive A/E pathogens (EPEC/EHEC) and SPI-1 *Salmonella* Typhimurium. Meanwhile, extraintestinal strains lacking a LEE island (UPEC, MG1655) showed non-significant partial growth inhibition (**Figure 2C–G**). This partial growth suppression in T3SS-deficient strains provided evidence that clofarabine entry is independent of the T3SS needle complex itself. As a purine nucleoside analogue, clofarabine likely acts as a prodrug requiring bacterial uptake via endogenous nucleoside transport pathways and intracellular phosphorylation by salvage enzymes [29]. [29]. In EPEC (and susceptible strains), these salvage kinases identify clofarabine as a false substrate, this accumulation disrupts nucleotide pool balance, which induces a state of active purine starvation [30, 31]. Interestingly, *Citrobacter rodentium* resistance to clofarabine sheds light into the potential mechanism of action of clofarabine through metabolic vulnerability. As an obligate mouse pathogen adapted to the highly inflamed nutrient-restricted colonic mucosa, C. *rodentium* possesses highly redundant nucleotide synthesis and salvage machinery capable of sustaining replication under severe metabolic stress [32–34]. While EPEC carries a similar machinery, it lacks the environmental priming to nucleoside stress of *C. rodentium* and is therefore more easily starved of dNTPs.

Finally, mass spectrometry analysis revealed a notable bacterial clearance mechanism that provided an explanation for the transient nature of clofarabine’s bacteriostatic effect (**Figure 2B**, **6F**, **6G**). Quantitative LC-MS analysis established that EPEC actively internalises intact clofarabine and enzymatically hydrolyses it into 2-deoxy-2-fluro-D-arabinose and 2-chloroadenine (**Figure 6F**, **6G**), indicative of N-glycosidic bond cleavage. This degradation is accompanied by a transcriptomic shift in purine catabolism and salvage pathway **(Figure 3E)**. This suggests that under clofarabine exposure, EPEC enters an “energy saving mode” that prioritises purine recycling to sustain basic metabolic survival; in doing so, this activated salvage machinery possess clofarabine as a false substrate, driving its enzymatic cleavage and ultimately enabling the bacteria to clear the drug and resume growth.

Beyond its bacterial mode of action, our evaluation of host cellular responses revealed that clofarabine appears to impact eukaryotic protein synthesis **(Figure 5D).** While clofarabine’s classic oncology mechanism focuses on DNA polymerases and ribonucleotide reductase inhibition, this host translational suppression would represent a previously unreported cellular effect. This dual host–pathogen translational impact emphasises the importance of host-side profiling when evaluating repurposed antimetabolites, providing key functional parameters for future drug optimisation.

In conclusion, this study highlights purine metabolic pathways as vulnerable and druggable targets for anti-virulence intervention in enteric pathogens. By establishing a mechanism by which clofarabine blocks T3SS effector translocation through translational suppression and purine competition, our study provides a proof-of-concept for repurposing nucleoside analogues to disarm bacterial infection machinery. While we also reveal an uncharacterised host translational suppression induced by clofarabine, these insights lay the groundwork for future medicinal chemistry studies to structurally modify clofarabine, optimising its bacterial sensitivity while eliminating its host translational suppression and systemic host toxicity. Furthermore, this work goes beyond clofarabine itself: the automated high-content screening pipeline established here provides a framework to evaluate broader chemical libraries and allows for the identification of novel anti-virulence hits.

## Methods

### Chemicals and Reagents

Clofarabine (ClF; Selleck Chemicals S1218) stock solutions were prepared at 10mM in dimethyl sulfoxide (DMSO; Sigma-Aldrich, D2438). Kanamycin (50ug/ml), ciprofloxacin (1ug/ml) and DMSO (1%) served as positive and negative controls, respectively. Standard reagents included LB broth (Sigma-Aldrich, L3522), LB agar (Sigma-Aldrich, L2897), Dulbecco’s modified Eagle Medium (DMEM; Themo Fisher Scientific, 11995065), L-glutamine (Themo Fisher Scientific, 25030149) and Penicillin-streptomycin (Thermo Fisher Scientific, 15140122).

### Bacterial Strains, Plasmids, and Growth Conditions

Bacterial strains and plasmids used in this study include, *E. coli* EPEC E2348/69 (wild-type O127:H6), EHEC EDL933 (Wild-type O157:H7), UPEC 125544 (Wild-type ExPEC), MG1655 (K-12 Wild-type), EPEC *ΔescN* (E2348/69 derivative), EPEC *ΔespG* (E2348/69 derivative), *Salmonella* Typhimurium SL1344 (peGFP-PltBA CdtB-pFPV mCherry) and *Citrobacter rodentium* DBS100 (ATCC 51459). Plasmids used in this study include pGEX2T-GFP (tac promoter-driven GFP expression, ampicillin resistant).

### Bacterial Growth and Drug Phenotyping Assays

For growth kinetics experiments, overnight EPEC cultures were inoculated into DMEM supplemented with FBS and HEPES to an initial OD600 of 0.1. Cultures were treated with CIF (10uM) or vehicle control and incubated at 37°C (200rpm). Optical density (OD600) was recorded every 30 min for 4.5h via spectrophotometry, or for 24h using a Tecan Sunrise microplane reader (Tecan, 30190079) in flat bottom 96-well plates (Corning, 3595). Minimum inhibitory concentration (MIC) determinations (0.1-10μM) and drug stability testing (pre-incubating CIF for 24h prior to inoculation) followed identical parameters.

### Mammalian Cell Culture and Infection Assays

HT1080 human fibrosarcoma epithelial cells (ATCC CCL-121) were maintained in DMEM supplemented with 10% FBS, 5% L-glutamine, 5% penicillin-streptomycin and 50ug/ml kanamycin at 37°C in a humidified 5% CO2 incubator. Cells were passaged every 3 days.

For infection assays, HT1080 cells (10^4^ cells/well) were seeded onto glass coverslips in 24-well plates and grown overnight. Logarithmic EPEC cultures were prepared by diluting overnight cultures 1:100 in fresh LB, growing to an OD600 of 1.0 then resuspended in supplement-free DMEM. Host cells were washed in the DMEM media and pre-treated with ClF (0.1-10μM) for 30 minutes and infected with EPEC at multiplicity of infection of 53. Plates were centrifuged at 1000 x*g* for 3 minutes and incubated for 3h at 37°C, 5% CO2.

### Immunofluorescence and microscopy

Fixed cells were blocked (3% BSA/ 0.2% Triton X-100), incubated with primary antibodies (anti-intimin 1:2000; anti-γH2AX 1:1000) for 1hr, followed by Alexa Fluor secondary antibodies (1:500) for 30minutes. Actin and host DNA synthesis were stained using Texas Red-X Phalloidin (1:250) and Click-iT EdU Alexa Fluor 647, respectively. Slides were mounted on Vectashield with DAPI and imaged on Nikon Ti Eclipse Widefield or Nikon A1 Confocal microscope and analysed in Fiji using previously published Fiji macros to automate the process.

### Secretome, Translation, and Immunoblotting

Whole cell lysates were collected, spun down and resuspended in SDS-Urea buffer. Culture secretome were clarified by spinning at 10000x*g* for 20 minutes, filter sterilised, TCA-precipitated in 20% TCA 1:1 dilution (final TCA concentration is 10%) overnight at 4°C, spun down at 10000x*g* for 20 minutes. Pellet is then washed in acetone and resuspended in SDS-Urea buffer 10x concentration relative to lysates. For IPTG induction, EPEC pGEX2T-GFP was induced with 0.5mM IPTG for 3hrs with hourly sampling.

Immunoblotting was carried out by resolving samples in 12%Bis-Tris-SDS-PAGE, stained with colloidal Coomassie or transferred to 0.2µm PVDF membrane (2.5 A,7minutes). Membranes were blocked (TBS 0.1% Tween20, 5% milk), incubated with primary antibodies at 1:1000 dilution: anti-Tir and anti-EspB (Gift from Prof. Vassilis Koronakis, University of Cambrdige), anti-BfpA (Gift from Prof. Gadi Frankel, Imperial College London), anti-GroEL (Sigma; #G6532) and anti-DnaK (Novus Biologicals; #NBP1-97490) then IRDye® 800CW Donkey anti-Mouse IgG (LiCor Biosciences; #926-32212) or IRDye® 680RD Donkey anti-Rabbit IgG (LiCor Biosciences; #925-68073) before imaging on the LI-COR Odyssey Sa.

### Bacterial RNA Isolation and Transcriptomic Profiling (RNA-seq)

Logarithmic EPEC E2348/69 cultures grown in supplemented DMEM were treated with clofarabine (10 *μ*M) or vehicle control. Samples were collected at specified time points, mixed immediately with RNAprotect Bacteria Reagent (Qiagen, 76506), and harvested by centrifugation. Total RNA was extracted using the RNeasy Mini Kit (Qiagen, 74104) with on-column DNase I digestion according to the manufacturer’s instructions. RNA concentration and integrity were evaluated using a NanoDrop spectrophotometer and Agilent 2100 Bioanalyzer. Ribosomal RNA was depleted using the Ribo-Zero rRNA Removal Kit (Illumina). cDNA libraries were constructed and sequenced on an Illumina platform generating 150 bp paired-end reads. Raw reads were quality-checked (FastQC), trimmed, and aligned to the EPEC E2348/69 reference genome. Differential gene expression analysis was conducted using DESeq2 in R, applying adjusted p-value thresholds (FDR < 0.05) and log_2_ fold-change cutoffs. The RNA-seq data generated in this study have been deposited in the NCBI Gene Expression Omnibus (GEO) database under accession code GSE346981.

### Mammalian Host Translation Assay (SUnSET Assay)

THP-1 human monocytes (ATCC TIB-202) were maintained in RPMI-1640 medium supplemented with 10% FBS, 2 mM L-glutamine, and 1% penicillin-streptomycin at 37°C in a humidified 5% CO2 incubator. For translational rate measurements via the Surface Sensing of Translation (SUnSET) assay, THP-1 cells (5 × 10^5^ cells/well in 24-well plates) were treated with increasing concentrations of clofarabine (0.1,1,10 *μ*M) or vehicle control for 3 h. During the final 30 min of incubation, puromycin dihydrochloride (Sigma-Aldrich, P8833) was added to the culture medium at a final concentration of 1 *μ*g/mL to label nascent polypeptide chains. Cells were harvested, washed in ice-cold PBS, and lysed in SDS-Urea lysis buffer. Protein samples were resolved by 12% Bis-Tris SDS-PAGE, transferred to 0.2 *μ*m PVDF membranes, and probed with anti-puromycin monoclonal antibody (1:1000; clone 12D10, Millipore #MABE343) followed by IRDye 800CW secondary antibody (1:10,000) for quantitative visualization on the LI-COR Odyssey Sa system.

### Metabolic Rescue and SOS Response Assays

For competitive metabolic rescue assays, logarithmic EPEC cultures in DMEM were co-treated with clofarabine (10 *μ*M) and increasing concentrations of exogenous adenosine (Sigma-Aldrich, A9251; 0-100 *μ*M). Bacterial growth was monitored continuously over 24 h via OD_#$$_ measurements, and extracellular secretome samples were collected and analyzed for Tir and EspB secretion by TCA precipitation and immunoblotting as described above. Induction of the bacterial SOS response was evaluated by monitoring RecA protein levels in EPEC lysates treated with clofarabine (10 *μ*M), ciprofloxacin (1 *μ*g/mL; positive control), or vehicle control via immunoblotting using anti-RecA primary antibody (1:1000; Abcam #ab63797).

### Mass Spectrometry (MS) Based Analysis

#### Analysis of EPEC Proteins by MS

Extracted proteins were separated by one-dimensional SDS-PAGE using a 12% polyacrylamide gel and visualised by Coomassie Blue staining. Gel bands of interest were excised and destained prior to in-gel digestion. Proteins were reduced with 50 mM tris(2-carboxyethyl)phosphine (TCEP) by incubation at 70°C for 20 min, followed by alkylation with 50 mM iodoacetamide (IAA) for 30 min at room temperature in the dark. Proteins were subsequently digested with trypsin overnight at a trypsin-to-protein ratio of 1:50 (w/w). Peptides were extracted from the gel pieces, collected, and dried using a vacuum concentrator. The dried peptide samples were resuspended in 60 µL of 0.5% (v/v) formic acid, of which 5 µL was used for subsequent LC–MS/MS analysis.

All mass spectrometry-based analyses were performed at the bioMICS Mass Spectrometry Facility, University of Sheffield, using an Orbitrap Exploris 480 mass spectrometer (Thermo Fisher Scientific) equipped with a nanospray ionisation (NSI) source and coupled to a Vanquish HPLC system (Thermo Fisher Scientific). Peptides were desalted online using a nano-trap column (75 μm i.d. × 20 mm; Thermo Fisher Scientific) and subsequently separated using an EASY-Spray column (50 cm × 50 μm i.d., PepMap C18, 2 μm particle size, 100 Å pore size; Thermo Fisher Scientific). Peptide separation was performed using a 100-min gradient with buffer B consisting of 0.1% (v/v) formic acid in 80% (v/v) acetonitrile. The gradient increased from 3% to 20% buffer B over 68 min, followed by an increase to 35% buffer B over 23 min, and to 99% buffer B over 1 min. The column was subsequently maintained at 99% buffer B for 9 min.

The mass spectrometer was operated in positive-ion mode using a data-dependent acquisition (DDA) method. The nanospray voltage was set to 1.7 kV and the ion-transfer-tube temperature to 275°C. Full MS spectra were acquired in the Orbitrap over an m/z range of 375–1200 at a resolution of 120,000, with a normalised automatic gain control (AGC) target of 300% (absolute AGC target, 3 × 10⁶), automatic maximum injection time and one microscan. Precursor ions exceeding an intensity threshold of 1 × 10⁴ and with charge states of 2–5 were selected for HCD fragmentation and up to 20 precursor ions were selected for fragmentation per acquisition cycle. Dynamic exclusion was set to 45 s after a single selection, with a ±10 ppm mass tolerance, and isotope exclusion was enabled. Precursors were isolated using a 2 m/z isolation window and fragmented by higher-energy collisional dissociation (HCD) at a normalised collision energy of 30%. MS/MS spectra were acquired in the Orbitrap at a resolution of 15,000, with the AGC target set to the instrument standard setting, automatic maximum injection time and one microscan. MS/MS spectra were acquired in centroid mode with the first mass set to m/z 120.

Raw mass spectrometry data were processed using MaxQuant (version 1.6.10.43). MS/MS spectra were searched against *E.coli* O127:H6 UniProt database (https://www.uniprot.org/). Database searches were performed using Trypsin/P as the proteolytic enzyme, allowing up to two missed cleavages. Carbamidomethylation of cysteine residues was specified as a fixed modification, whereas oxidation of methionine and protein N-terminal acetylation were specified as variable modifications. Peptide and protein identifications were controlled at a false discovery rate (FDR) of 1% using the target-decoy strategy implemented in MaxQuant. FDR thresholds of 0.01 were applied at both the peptide and protein levels.

### Analysis of Clofarabine and its Degradation by MS

#### Bacterial Culture and Sample Preparation

Enteropathogenic *E. coli* (EPEC) cultures were grown in Dulbecco’s Modified Eagle Medium (DMEM) supplemented with 10% fetal bovine serum (FBS), in the presence or absence of 10 µM clofarabine. Samples were collected at optical densities (OD₆₀₀) of 0.1, 0.2, and 1.0. Bacterial cultures were centrifuged at 3,000 × g for 10 min to separate bacterial cells from the extracellular fraction.

For analysis of extracellular clofarabine and its putative degradation product, culture supernatants were filter-sterilised using a 0.45 µm filter. Proteins were precipitated by addition of four volumes of acetone (50 µL sample and 200 µL acetone; 1:4, v/v), followed by incubation at −20°C overnight. Samples were subsequently centrifuged at 20,000 × g for 10 min, and the resulting supernatants containing the soluble small-molecule fraction were collected for LC–MS/MS analysis.

For analysis of intracellular clofarabine and its putative degradation product, bacterial pellets were washed with cold water and centrifuged at 7,000 × g for 2 min. The washed bacterial pellets were resuspended in 100 µL methanol and lysed using an electric homogeniser. Cell lysates were centrifuged at 20,000 × g for 10 min to remove insoluble cellular debris, and the resulting supernatants containing the extracted intracellular metabolites were collected for LC–MS/MS analysis.

#### LC–MS/MS Analysis of Clofarabine and Its Degradation Product

Extracellular and intracellular extracts were analysed using the same HPLC and MS system as described above. A 1 μL aliquot of each sample was injected for LC–MS/MS analysis. Chromatographic separation was performed in a trap-and-elute configuration using a 300 μm × 0.5 cm trap column and an ES903 separation column (75 μm i.d. × 50 cm; Thermo Fisher Scientific). Mobile phase A consisted of water containing 0.1% (v/v) formic acid, and mobile phase B consisted of 80% (v/v) acetonitrile containing 0.1% (v/v) formic acid. The flow rate was maintained at 250 nL/min. The gradient commenced at 10% B and increased to 90% B over 15 min, followed by an increase to 100% B by 18 min. The column was maintained at 100% B until 25 min for column washing, followed by column re-equilibration.

The mass spectrometer was operated in positive-ion mode using a data-dependent acquisition (DDA) method optimised for small-molecule analysis. The nanospray voltage was set to 1.7 kV and the ion-transfer-tube temperature to 275°C. Full MS spectra were acquired in the Orbitrap over an m/z range of 70–800 at a resolution of 60,000. The automatic gain control (AGC) target was set to the instrument standard setting, with a maximum injection time of 100 ms and one microscan. Full MS spectra were acquired in profile mode. For data-dependent MS/MS acquisition, precursor ions exceeding an intensity threshold of 5 × 10³ were selected for fragmentation, with up to four dependent MS/MS scans acquired per acquisition cycle. Dynamic exclusion was applied after one precursor selection for 2.5 s using a ±5 ppm mass tolerance, with isotope exclusion enabled. Precursor ions were isolated using a 1.5 m/z isolation window and fragmented by higher-energy collisional dissociation (HCD) at a normalised collision energy of 35%. MS/MS spectra were acquired in the Orbitrap at a resolution of 15,000, with the AGC target set to the instrument standard setting, a maximum injection time of 22 ms, and one microscan. MS/MS spectra were acquired in profile mode.

Raw LC–MS data were analysed using FreeStyle software (Thermo Fisher Scientific). Extracted-ion chromatograms (XICs) were generated using a mass tolerance of ±5 ppm. Clofarabine was monitored at m/z 304.06, whereas its putative degradation product (fluorinated purine) was monitored at m/z 170.02. For quantitative analysis of clofarabine, an external calibration curve was generated using clofarabine standards at concentrations of 1 nM, 10 nM, 100 nM, and 1 μM. Extracted-ion chromatographic peak areas were integrated using FreeStyle software and plotted against clofarabine concentration. Linear regression was used to generate the calibration curve, which showed good linearity over the concentration range examined (R² = 0.9974). Clofarabine levels in experimental samples were determined by comparison of their extracted-ion chromatographic peak areas with the calibration curve.

In the absence of an authentic standard for the m/z 170.02 species, the fluorinated purine product was assessed semi-quantitatively relative to clofarabine. The extracted-ion chromatographic peak area at m/z 170.02 was therefore compared with that of clofarabine (m/z 304.06), rather than being used to calculate an absolute concentration. The assignment of the m/z 170.02 species as a putative clofarabine degradation product (fluorinated purine) was based on its detection following bacterial exposure and the expected cleavage of the nucleoside glycosidic bond. This interpretation is consistent with previous observations that bacteria can metabolise structurally related halogenated purine nucleosides through cleavage of the glycosidic bond to generate the corresponding purine-base products [35].

### Statistical Analysis

Data are expressed as mean ± SEM of at least three independent biological replicates (*n* ≥ 3). Statistical analyses were performed using GraphPad Prism (version 9.0; GraphPad Software, San Diego, CA, USA) and RStudio (version 2026.01.0+392).

