## Supplementary Figure 1 for "Repurposing clofarabine against enteric pathogens via metabolic starvation"

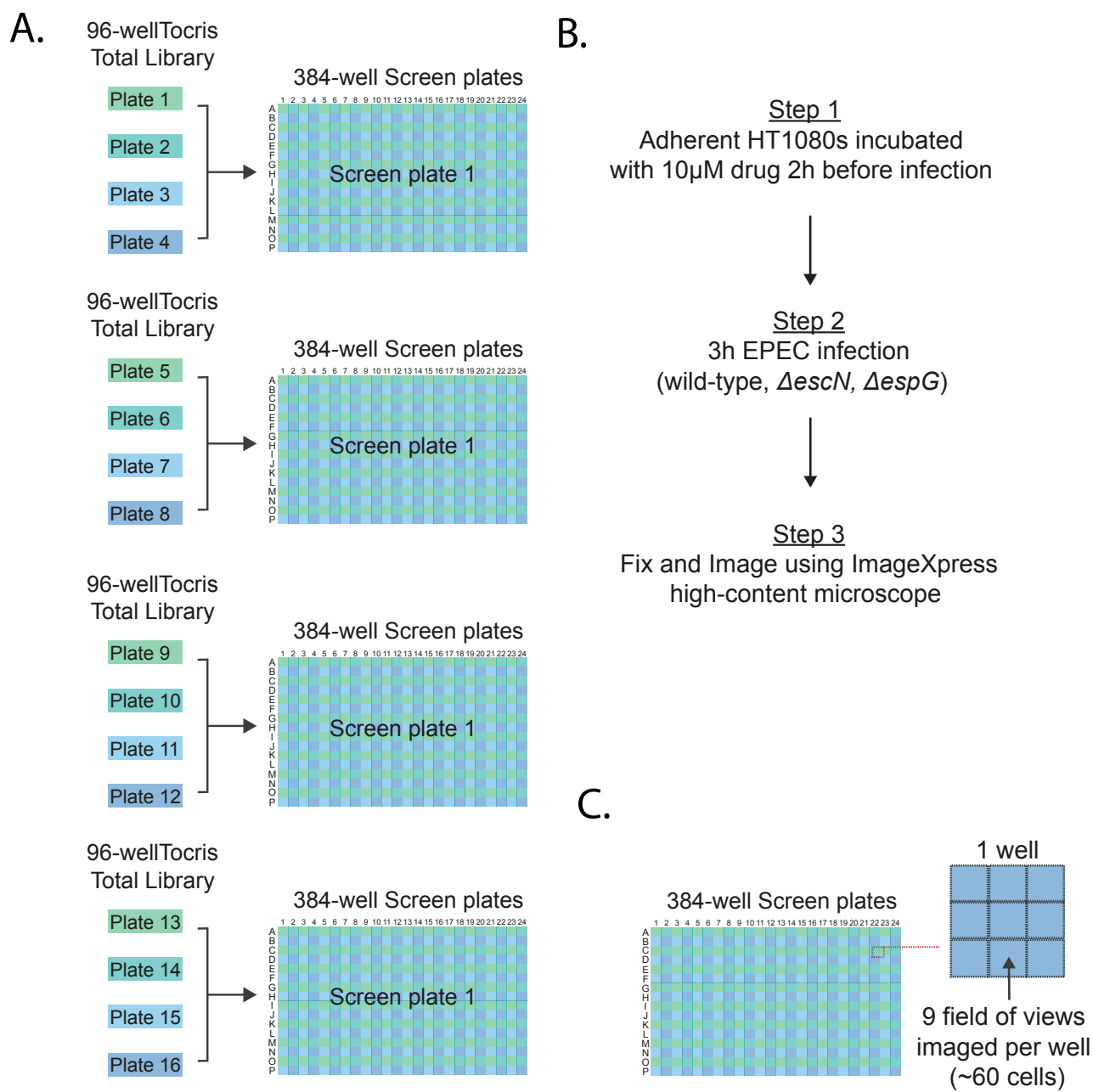

**Figure S1: High-content screening workflow for identifying small-molecule inhibitors of EPEC pedestal formation.** (A) Schematic representation of drug library transfer and plating format. Compounds from the Tocriscreen Total Library (1,120 molecules in 96-well master plates) were arrayed into 384-well assay plates. Each compound was screened in quadruplicate (4 technical replicates) across 2 independent biological experiments. (B) Experimental workflow of the automated high-content screen. Adherent HT1080 host cells were pre-treated with 10 μM compound for 2 h prior to infection with wild-type (WT) EPEC, $\Delta escN$ (T3SS-deficient control), or $\Delta espG$ mutant strains for 3 h. Plates were subsequently fixed, fluorescently stained, and imaged using an ImageXpress high-content microscope. (C) Imaging strategy per well. Nine independent fields of view (~60 host cells per well) were automatically acquired and analysed per condition to quantify actin pedestal formation.
