## Supplementary figures and images for "Repurposing clofarabine against enteric pathogens via metabolic starvation"

### Supplementary Figure 2

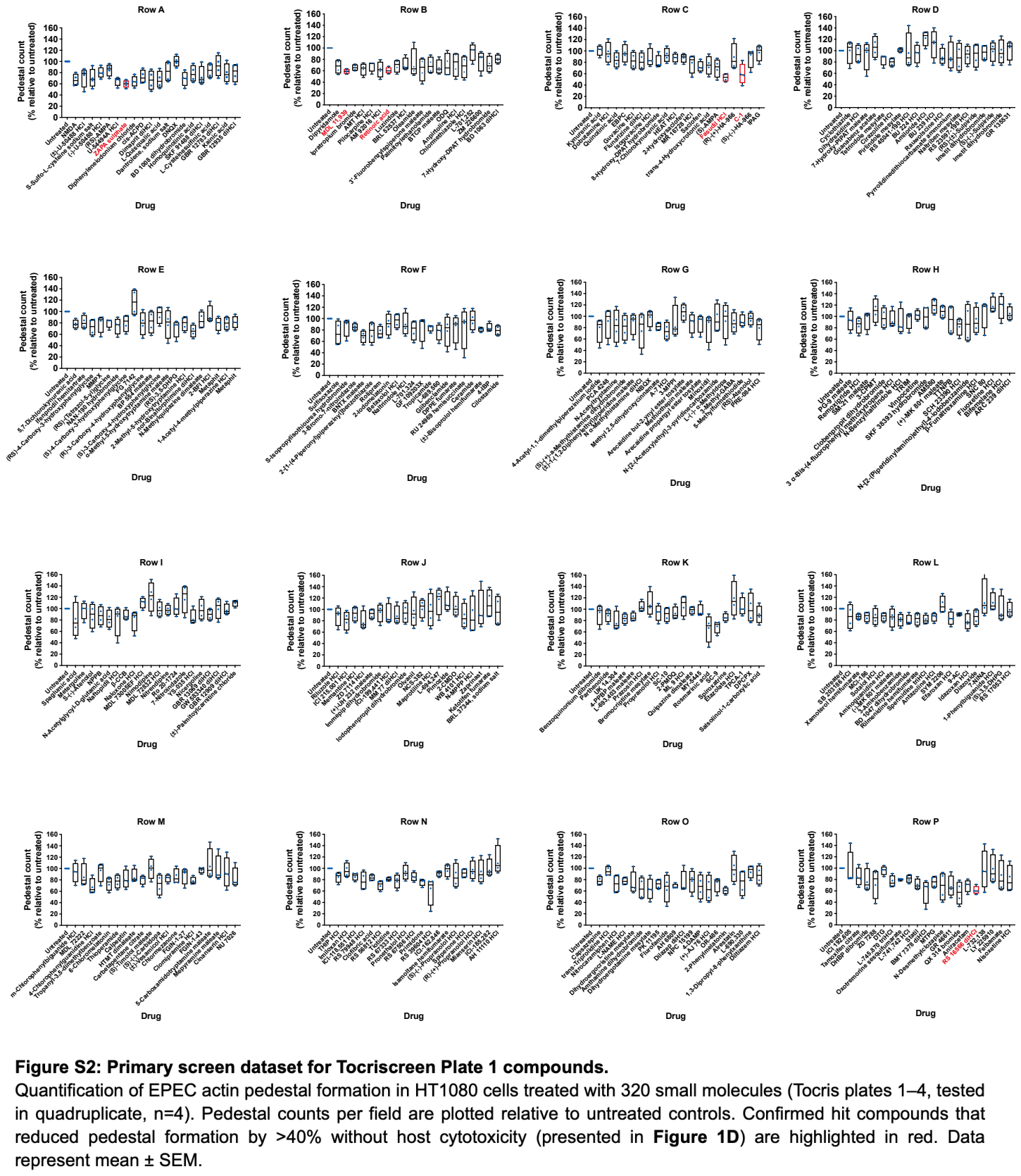

### Supplementary Figure 4

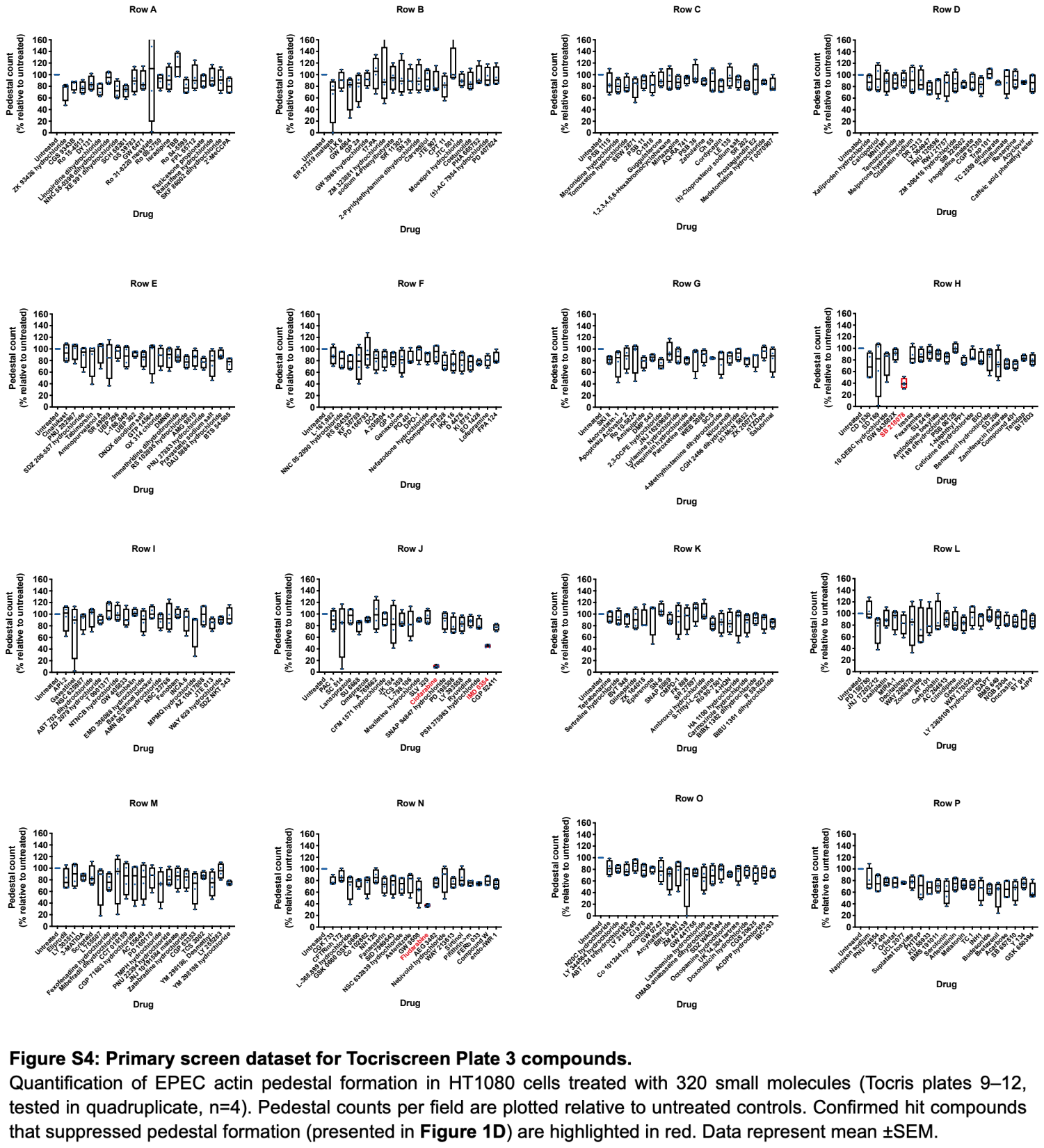

### Supplementary Figure 5

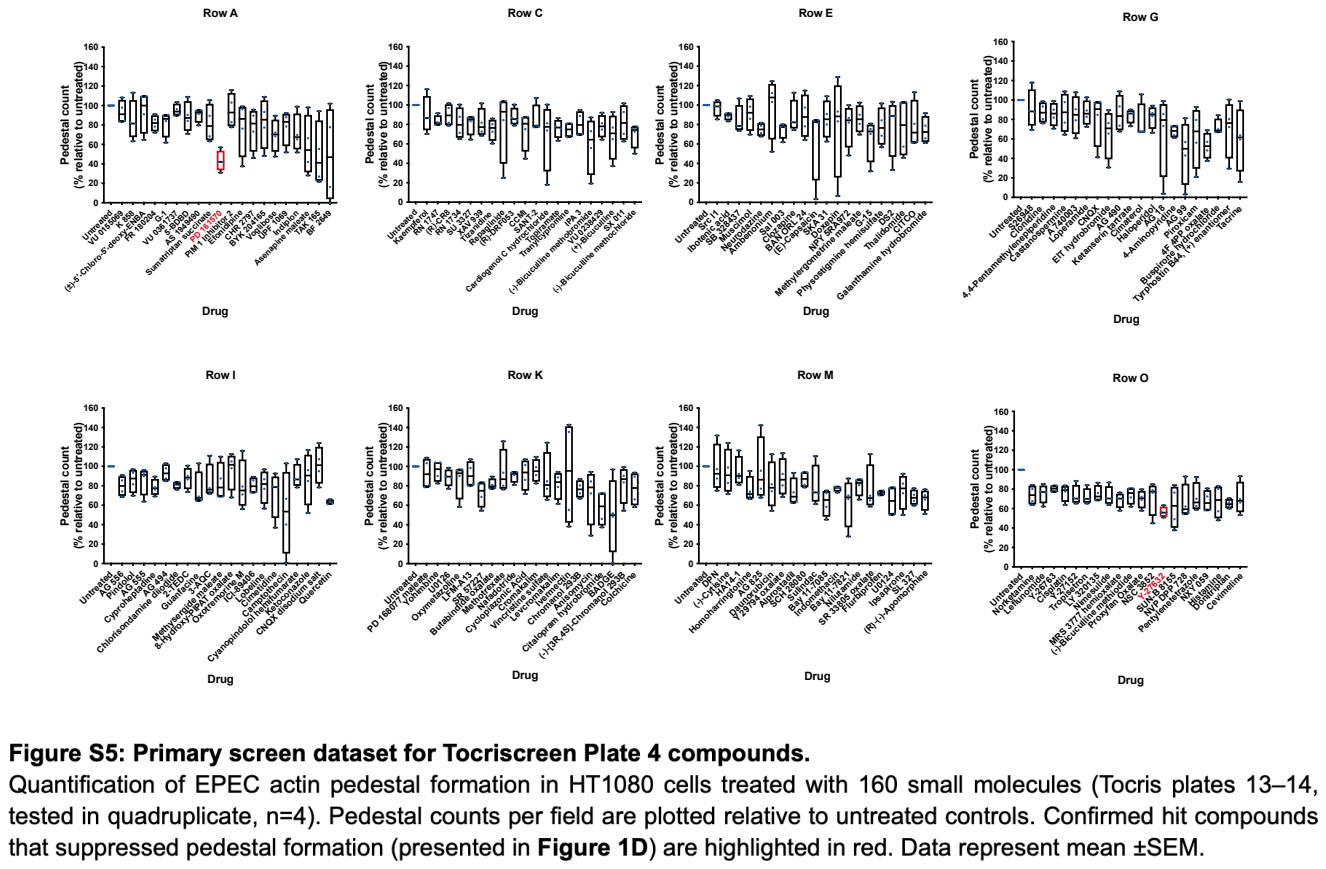
