## Supplementary Figure 3 for "Repurposing clofarabine against enteric pathogens via metabolic starvation"

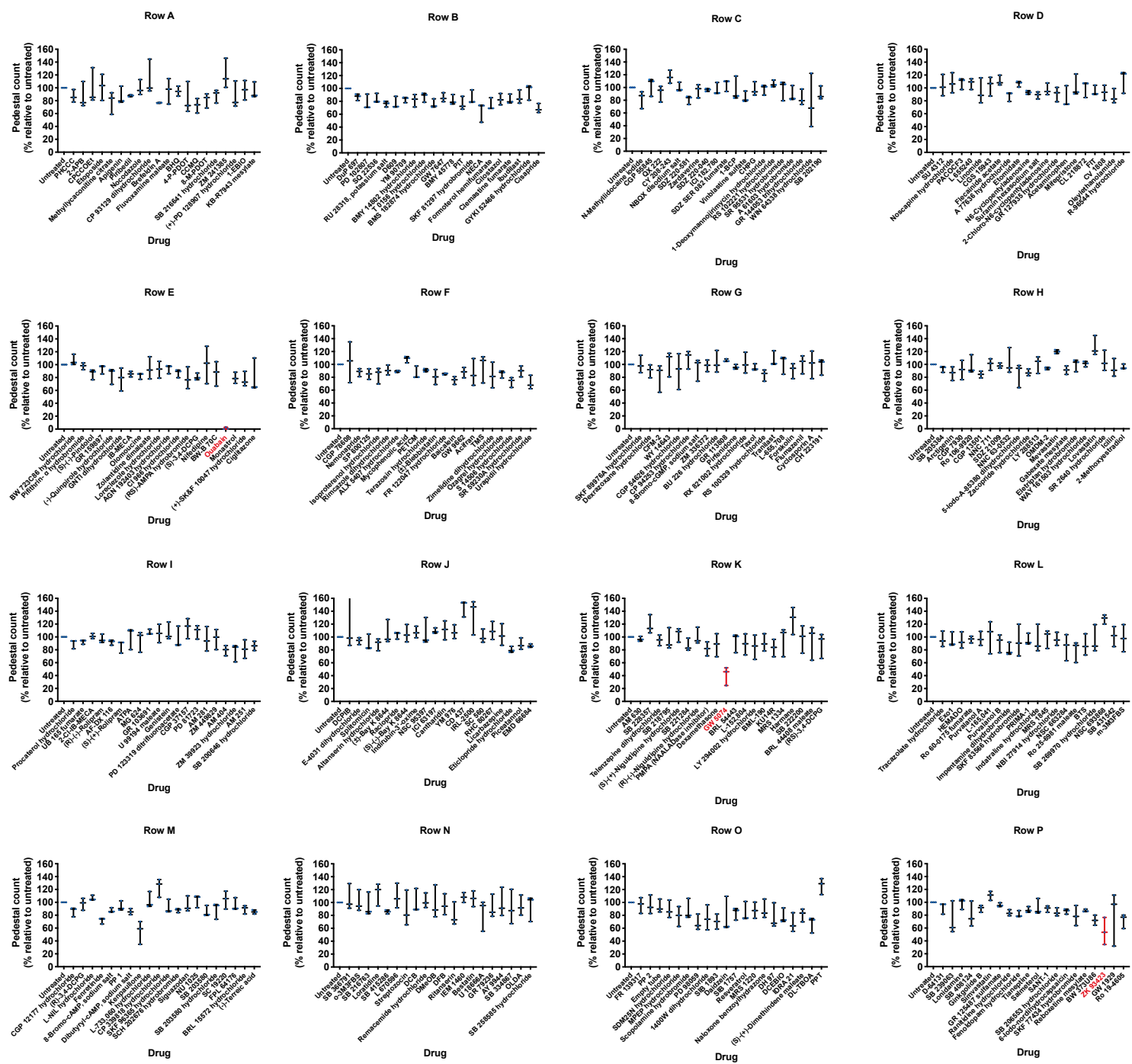

**Figure S3: Primary screen dataset for Tocriscreen Plate 2 compounds.**

Quantification of EPEC actin pedestal formation in HT1080 cells treated with 320 small molecules (Tocris plates 5–8, tested in quadruplicate, n=3). Pedestal counts per field are plotted relative to untreated controls. Confirmed hit compounds that suppressed pedestal formation (presented in **Figure 1D**) are highlighted in red. Data represent mean ± SEM.
