## Supplementary Figure 6 for "Repurposing clofarabine against enteric pathogens via metabolic starvation"

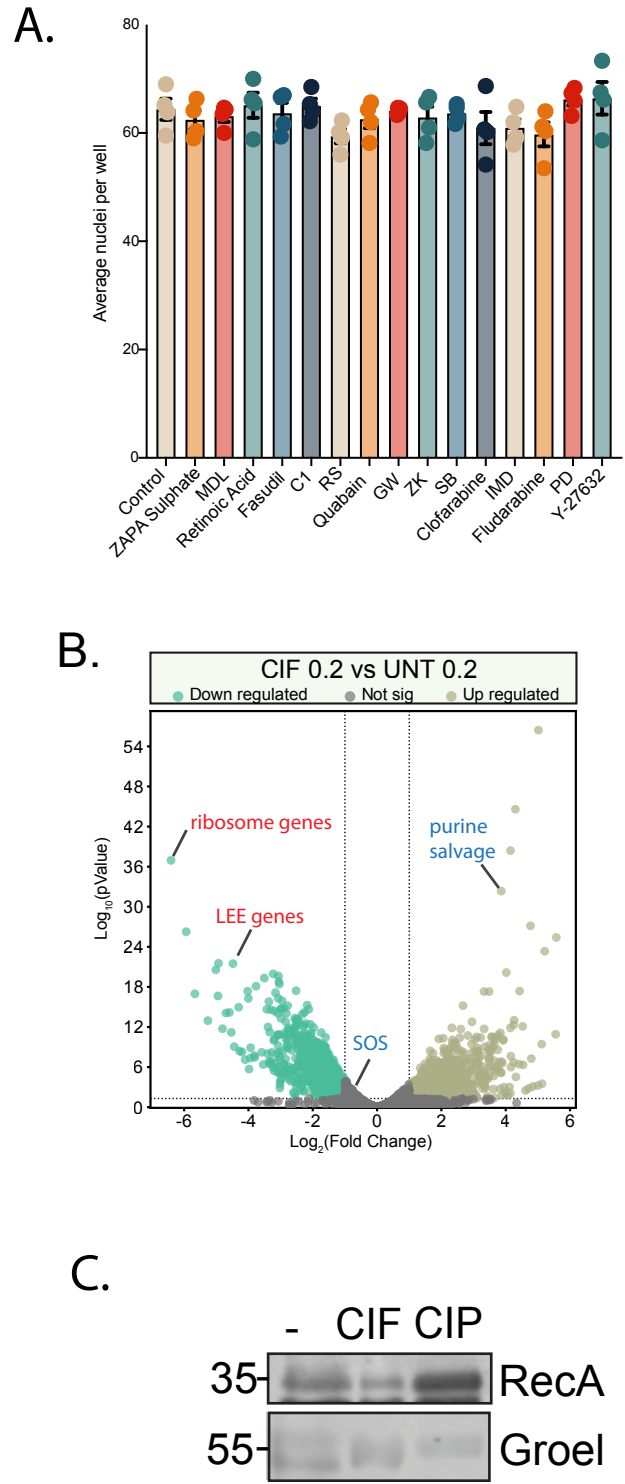

**Figure S6: Host cell viability, global transcriptional response, and DNA damage assessment under clofarabine treatment.**

**(A)** Host cell nuclei counts following treatment with top hit compounds from **Figure 1D**. HT1080 host cell nuclei per well were quantified, demonstrating that the observed reduction in EPEC actin pedestals is not due to host cell toxicity or detachment. **(B)** Volcano plot illustrating differentially expressed EPEC genes upon 10 μM clofarabine (CIF) treatment at OD600 =0.2 relative to untreated control (UNT), supporting **Figure 3B**. Downregulated gene clusters (green), including examples for locus of enterocyte effacement (LEE) and structural ribosomal operons, and upregulated clusters (gold), including purine salvage pathways, are highlighted. SOS response genes remain unchanged. **(C)** Immunoblot analysis of RecA expression in EPEC whole-cell lysates treated with vehicle (-), clofarabine (CIF, 10 μM), or ciprofloxacin (CIP, positive control for SOS/DNA damage induction). GroEL was probed as a loading control.
