## Supplementary Figure 7 for "Repurposing clofarabine against enteric pathogens via metabolic starvation"

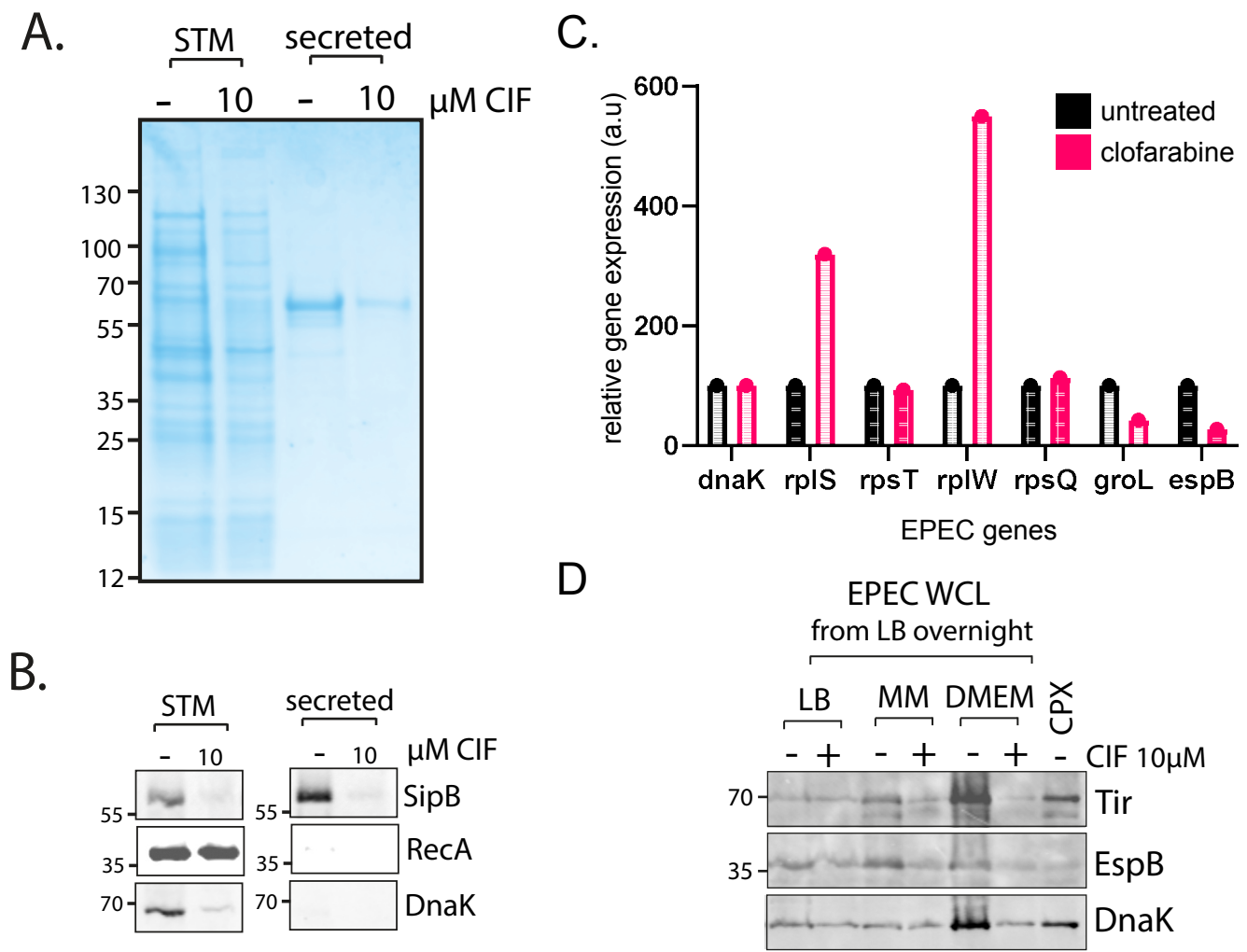

**Figure S7: Clofarabine-mediated suppression of *Salmonella* Typhimurium, ribosomal genes and nutrient-dependent reversal of clofarabine activity.**

**(A)** Coomassie blue-stained SDS-PAGE gel of whole-cell lysates (STM) and filtered culture supernatants (secreted) from *Salmonella* Typhimurium SL1344 grown in the presence (+) or absence (-) of 10 μM clofarabine (CIF). Total protein staining served as the primary loading control due to drug-induced repression of endogenous chaperones. **(B)** Immunoblot analysis of *Salmonella* whole-cell lysates and secretome probing the T3SS translocator SipB, RecA, and DnaK loading control, confirming clofarabine-mediated blockade of T3SS secretion in *Salmonella*. **(C)** qRT-PCR relative gene expression normalised to internal *dnaK* within each respective treatment condition (untreated vs. clofarabine-treated EPEC). Conventional self-normalisation creates an artificial apparent upregulation of target transcripts (*rplS*, *rplW*) because clofarabine concurrently represses the internal reference gene *dnaK*. **(D)** Immunoblot analysis of EPEC whole-cell lysates (WCL) cultured overnight in rich medium (LB), minimal medium (MM), or DMEM in the presence (+) or absence (-) of 10 μM clofarabine (CIF) or ciprofloxacin (CPX control). Clofarabine fails to repress Tir, EspB, and DnaK in rich LB media, demonstrating that abundant extracellular purines and metabolites out-compete clofarabine activity.
